# Avian interferon regulatory factor (IRF) family reunion: IRF3 and IRF9 found

**DOI:** 10.1101/2024.09.24.613690

**Authors:** Lenka Ungrová, Josef Geryk, Marina Kohn, Dana Kučerová, Veronika Krchlíková, Tomáš Hron, Vladimír Pečenka, Petr Pajer, Eliška Gáliková, Ľubomíra Pecnová, Bernd Kaspers, Jiří Hejnar, Jiří Nehyba, Daniel Elleder

## Abstract

Interferon regulatory factors (IRFs) are a family of transcription factors in jawed vertebrates with important functions in immunity and many other key cellular processes. The genomes of most vertebrates encode ten IRF genes (IRF1 to IRF10). IRF3 and IRF9 have key roles in the interferon (IFN) induction and signaling. Most of our knowledge about the IFN pathways originates from the study of the mammalian IFN system, and the description of the corresponding avian components is not as complete. Both IRF3 and IRF9 were considered missing from the chicken genome and also from the genomes of all other avian species. Here we describe multiple avian IRF3 and IRF9 genes, all with difficult GC-rich sequence context which prevented their earlier characterization. IRF3 orthologs are more narrowly distributed and are present in the avian infraclass Palaeognathae residing in a syntenic genomic locus shared with other vertebrates. In contrast, IRF9 orthologs were found in most avian species with the notable exception of the order Galliformes. In about half of the avian orders analyzed, IRF9 was located in noncanonical chromosomal positions indicating past evolutionary translocations. Importantly, phylogenetic analysis confirmed the correct orthology of all newly described avian IRFs. We performed a series of experiments using duck (*Anas platyrhynchos*) IRF9, confirming its key role in the IFN signaling pathway. Knockout of IRF9 in duck embryonal fibroblasts decreases the induction of IFN-stimulated genes (ISGs). Full induction of ISGs in duck cells requires both intact IRF9 and canonical IFN-stimulated response element (ISRE). Lastly, intact IRF9 is needed for IFN-mediated protection of duck cells against vesicular stomatitis virus (VSV)-induced cytopathic effect. The identification of avian IRFs fills an important gap in our understanding of avian immunology and brings new questions related to the evolution of the IRF family.

## INTRODUCTION

The interferons are key cytokines orchestrating the antiviral defense and immune regulation (García-Sastre and Biron 2006). Although interferon (IFN) was originally discovered during work with embryonated chicken eggs (Krause and Pestka 2007), most of the subsequent molecular dissection of IFN-related pathways was performed in mammalian cells (reviewed in (Stark 2007; Philips et al. 2022)). Conceptually, the first stage of these pathways involves IFN induction, where various sensor families detect viral products and through activation of several transcription factors induce production of these cytokines. Second stage is IFN signaling, during which secreted type I interferons including IFN-β and multiple IFN-α interact with type I IFN receptor (IFNAR) and initiate a cascade of events resulting eventually in phosphorylation of signal transducers and activators of transcription (STAT) molecules. Various STAT complexes translocate to the nucleus and bind to genetic regulatory elements like the IFN-stimulated response element (ISRE). This leads to the activation of hundreds of IFN-stimulated genes (ISGs) and establishment of an antiviral state (Schoggins 2019). Importantly, both IFN pathway stages operate using transcription factors of the interferon regulatory factor (IRF) family. Two IRF family members, IRF3 and IRF7, are activated by upstream kinases and induce promoters of IFN genes. Subsequently, during the IFN signaling stage, IRF9 forms a trimeric complex with STAT1 and STAT2, also called IFN-stimulated gene factor 3 (ISGF3), which binds to ISRE. In contrast to substantial knowledge about mammalian IFN system, the understanding of the avian counterparts of transducers of IFN signaling is not as complete. Mainly in the last two decades following the publication of the chicken genome sequence, many of the key components of the chicken innate immune defense have been described (Campbell and Magor 2020; Kaiser 2012). A striking observation is the apparent lack of identified IRF3 and IRF9 in avian species (Santhakumar et al. 2017; Magor et al. 2013).

In general, IRFs of jawed vertebrates (Gnathostomata) are a family of eleven genes that encode transcription factors with important functions in immunity, hematopoietic differentiation, and other cellular processes (Negishi, Taniguchi, and Yanai 2018). The IRF paralogs which number consecutively from IRF1 to IRF11 belong, according to their sequence similarities, to three gene groups: IRF1-G (which includes IRF1, 2, 11), IRF3+5-G (IRF3, 5, 6, 7), and IRF4-G (IRF4, 8, 9, 10) (Nehyba, Hrdlicková, and Bose 2009; Du et al. 2018; Shu, Sun, and Xu 2015). Synteny analysis (Fig. 1, Suppl. Tab. 1) suggests that each of the groups evolved in ancient vertebrates from a single gene predecessor by two subsequent whole genome duplications (2WGD) during the older paleozoic before the split of chondrichthyan and osteichthyan vertebrate lineages (Simakov et al. 2020). Additional IRF paralogs might have evolved later including those in vertebrate species that underwent a third or more successive genome duplications, like in teleost fish (Clark, Boudinot, and Collet 2021).

**Figure 1.**
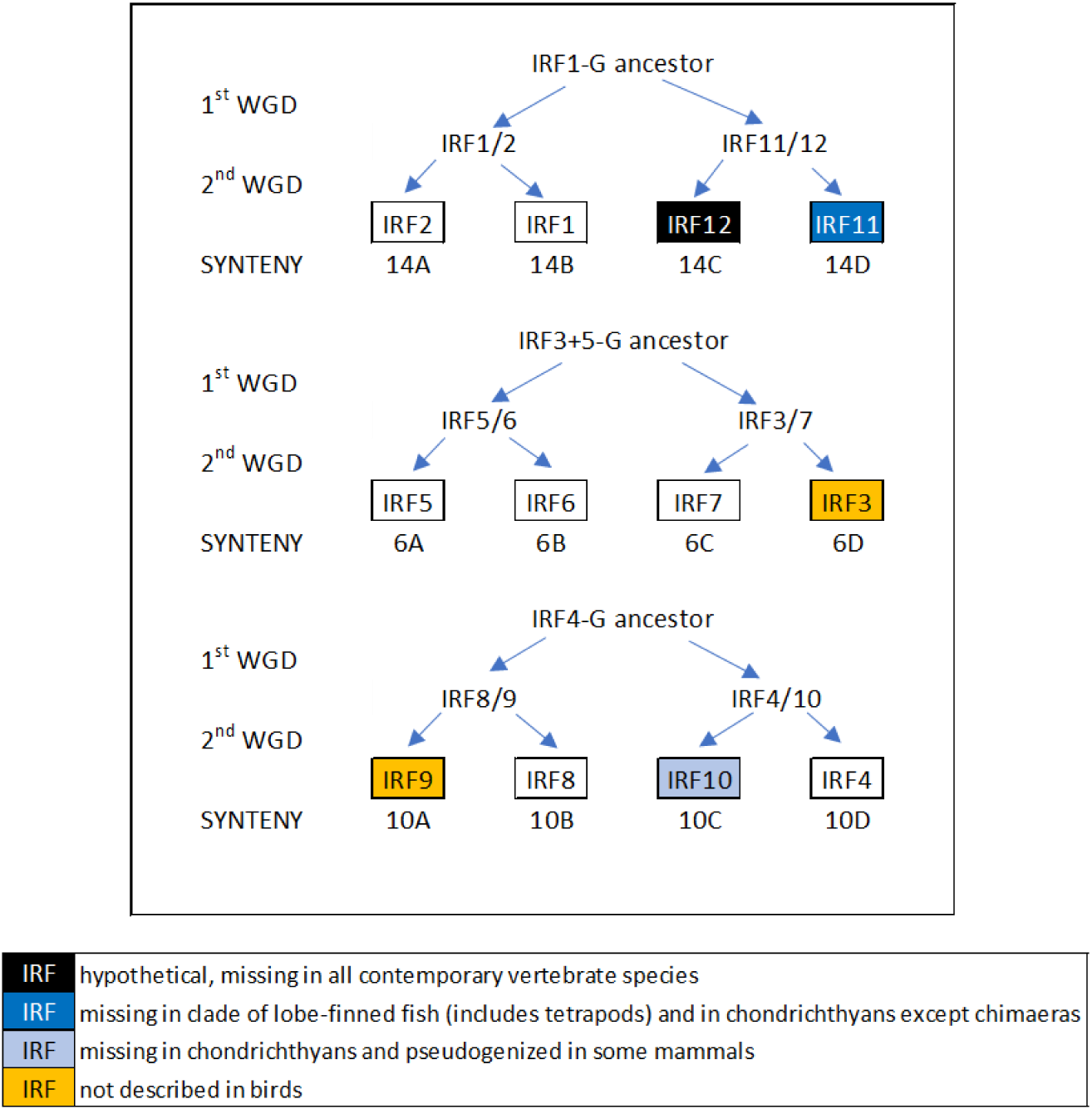
Synteny supports the hypothesis of the origin of the jawed vertebrate IRF gene family by two-fold genome duplication. The 2WGD event led to the quadruplication of 17 gene groups (paralogons), each representing one chromosome of the pre-WGD chordate, into four quads designated A – D resulting in 68 gene groups described and numbered from 1A to 17D by (Lamb 2021). Syntenic analysis indicates that IRF genes of each of the three IRF groups always belong to the same paralogon: paralogon 14 (IRF1-G: IRF1, 2, 11), paralogon 6 (IRF3+5-G: IRF3, 5, 6, 7), and paralogon 10 (IRF4-G: IRF4, 8, 9, 10), and at the same time always to a different quad (A, B, C, D), indicating an origin from a single prevertebrate chordate gene for each group. Due to subsequent paralog loss, not all genes are present in all four copies in contemporary species. IRF4-G genes were classified as pertinent to paralogon 10 by (Lamb 2021). For paralogon classification of IRF1-G and IRF3+5-G genes see Suppl. Tab. 1.

Unequal presence of IRF paralogs in different vertebrate species indicates that the 2WGD event has been followed by the loss of some IRF genes. Nevertheless, a large number of amniote species have ten genes, IRF1 to IRF10, and most of the observed paralog absences except IRF10 pseudogenization in some mammals are likely artifacts of incompleteness of genome sequences (Li et al. 2023). Therefore, the inability to discover IRF3 and IRF9 in avian species always seemed peculiar (Santhakumar et al. 2017). Some reports mentioned the presence of these two genes in the chicken but that was always the result of confusion with other IRF paralogs. IRF3 was often confused with IRF7 (see (Cheng et al. 2019)). The gene annotated as IRF9 in the chicken genome in GenBank and cited in some reports as such is actually the IRF10 gene described in the chicken previously (Nehyba et al. 2002).

We have previously shown that a subset of avian genes is encoded by sequences rich in G and C nucleotides (GC-rich). Such sequences are hard to identify by next generation sequencing and also difficult to amplify by PCR. Therefore, such genes were often falsely reported as missing from avian genomes (Hron et al. 2015; Huttener et al. 2021). We published and characterized several such “hidden” avian genes, including e.g. TNF-α, FoxP3, LAT and leptin (Rohde et al. 2018; Burkhardt et al. 2022; Janusova et al. 2023; Seroussi et al. 2019). In this work, we have focused on the discovery of avian IRF9 and IRF3. We report the identification of GC-rich IRF9 in multiple avian species, and present experimental characterization of duck (*Anas platyrhynchos*) IRF9. We also identified GC-rich IRF3 in paleognath birds.

## RESULTS

### 1) IRF3 genes were found only in genomes of basal avian infraclass of Palaeognathae

Systematic search in avian sequences deposited in National Center for Biotechnology Information (NCBI) public databases including high throughput sequencing repositories (Sequence Read Archive; SRA) resulted in the discovery of the first true avian IRF3 gene. The sequence of the gene found in nucleic acid data of emu (*Dromaius novaehollandiae*) was verified by de novo assembly from SRA RNASeq archives (Suppl. Tab. 2). The GC content of this sequence was high (77%). The emu IRF3 was compared with sequences of other emu IRF paralogs and with complete sets of IRF genes from additional two avian, two mammalian and one teleost species (Fig. 2). The tree also included newly discovered duck and emu IRF9 genes that will be discussed later. A phylogenetic tree showed the emu gene (black arrow) co-localized with the other previously characterized vertebrate IRF3 genes in one monophyletic group, confirming its IRF3 identity. The branching order of the tree was in general agreement with the syntenies shown in introductory Fig. 1 (except the branching of IRF8 and IRF9 as well as likely artificial clustering of trout IRF5 with IRF6 proteins) and also agreed with the IRF3 genes being the members of the higher order group, IRF3+5-G.

**Figure 2.**
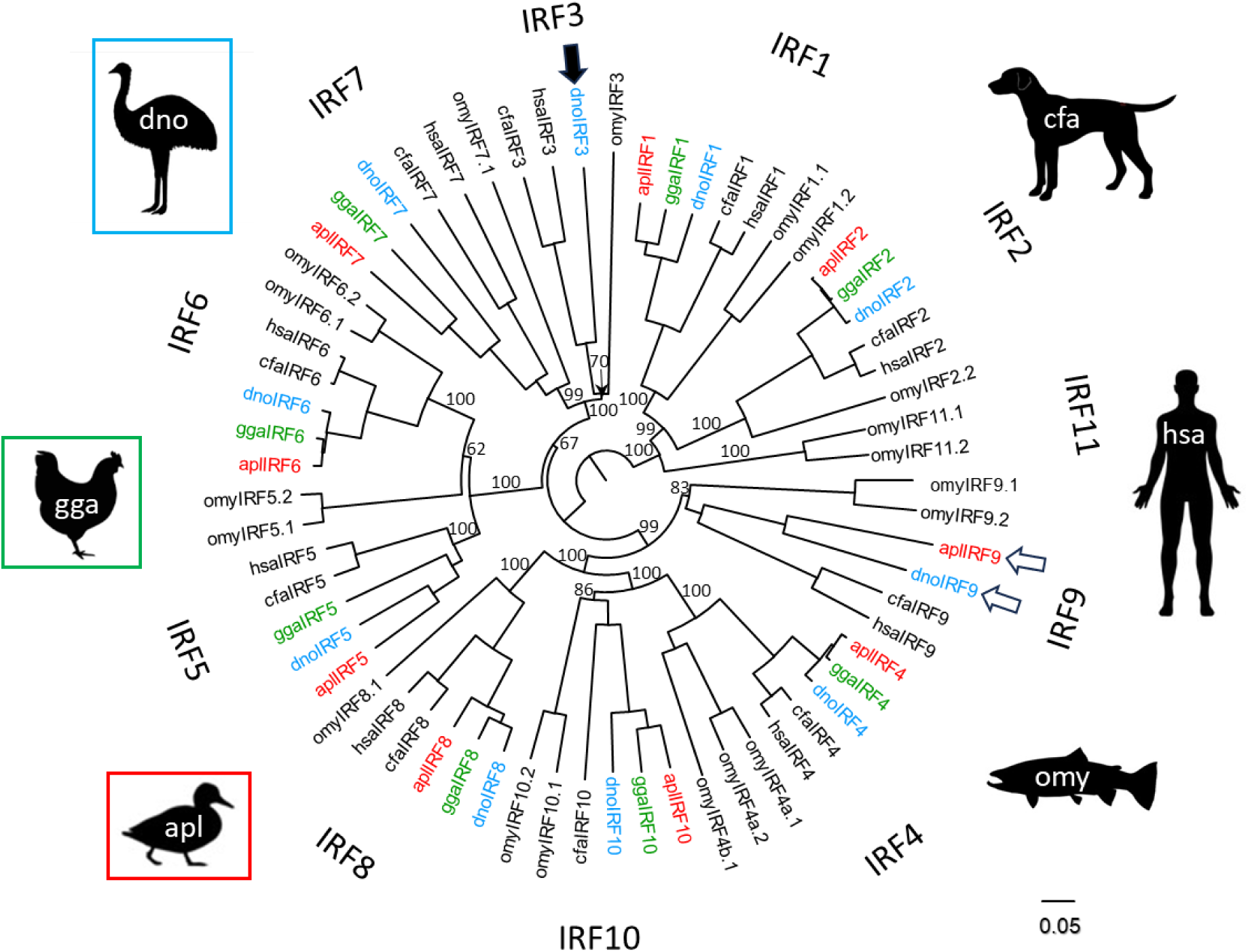
Position of avian IRF3 and IRF9 in the evolutionary tree of the IRF family. The black arrow and the white arrows indicate the respective positions of avian IRF3 and avian IRF9 genes described in this report. Protein sequences of all IRF genes in six vertebrate species: rainbow trout (*Oncorhynchus mykiss* – **omy**), emu (*Dromaius novaehollandiae* – **dno**), chicken (*Gallus gallus* – **gga**), duck (*Anas platyrhynchos* – **apl**), human (*Homo sapiens* – **hsa**), and dog (*Canis familiaris* – **cfa**), were compared. Fish family Salmonidae which includes rainbow trout has all 11 vertebrate IRF genes. Most of these IRF paralogs are present in multiple copies as a result of multiplication of the original 11 genes in additional teleost and salmonid specific whole genome duplications. The IRF11 gene wasn’t found in any tetrapod species. Humans lack functional IRF10 gene. In chicken no IRF3 or IRF9 genes were identified and are likely missing. Protein sequences were extracted from NCBI protein databases or were assembled/predicted (emu IRF3 and IRF5, duck IRF5 and IRF9) from SRA database reads and genome contig sequences (see details in Suppl. Tab. 2 and 5). The sequences were aligned by ClustalX, a neighbor-joining (NJ) tree constructed and visualized by FigTree. Branch support values calculated by bootstrapping (10000 replicates) are shown as percentages and only for central branches of the tree. The scale bar indicates the number of amino acid substitutions per site.

During the work on this manuscript a new emu genome was released (GCF_036370855.1) that contained the IRF3 gene predicted by NCBI annotation pipeline. This annotation mostly agrees with our coding sequence (CDS) except that the NCBI predicted genomic sequence has the N-terminal extension and it also contains what we consider to be an error – break in the reading frame in exon 6 and compensatory addition of one superfluous exon which is not supported by RNASeq data. Interestingly both our sequence and the genomic version contain in-frame extension of the second IRF3 exon by four-fold repetition of 12-20 amino acid codons at the 3’ boundary of the exon. The emu IRF3 gene is localized on microchromosome 31 and is surrounded by the genes belonging to group 6D of ohnologs (e.g. LRRC4B, MYBPC2, SCAF1, SHANK1, SYT3, TRIM28). This syntenic position which follows the original IRF3 synteny established after the second WGD provides additional verification that the gene is indeed IRF3.

The emu IRF3 gene sequence was used to search for IRF3 genes in other avian genomes and transcriptomic data. The search resulted in the assembly of one complete and one almost complete sequence from ostrich (*Struthio camelus*) and Darwin’s rhea (*Rhea pennata*), respectively (Suppl. Tab. 2, Suppl. Fig. 1). None of these two sequences contained above mentioned exon 2 in-frame extension indicating that this structural variation is likely limited to emu and possibly to emu-related species. The IRF3 sequences in all three birds were surprisingly similar (88%, 84%, and 85% of identity between emu-ostrich, emu-rhea, and rhea-ostrich) despite estimated divergence of these species before approximately 80 million years (Phillips et al. 2010). The GC content of all three sequences was in the range of 76-77%. Compared to the seven exon structure of mammalian IRF3 genes, avian genes have nine exons similar to genes of non-avian archosaurs (crocodiles + turtles). The genome contig containing *Rhea pennata* IRF3 sequences also contained MYBPC2 gene indicating that rhea IRF3 is likely in the same original syntenic position as in emu. Further blast searches in avian genome databases resulted in finding partial IRF3 sequences in two additional species of Palaeognaths (*Nothoprocta perdicaria*, *Tinamus major*)(Suppl. Tab. 2).

In contrast, no IRF3 was found in 8 high-quality genomes representing a wide range of neognath species (Suppl. Tab. 3). In all cases these neognath genomes contained SCAF1 gene which is located on the chromosome directly besides IRF3 in all genomes of non-avian jawed vertebrates we examined as well as in the genome of emu. Presence of SCAF1 suggests that neognath genomes used to search for IRF3 are not missing sequences of chromosomal regions where IRF3 would be expected to be located if conserved synteny would be preserved. Search in nr/nt genbank databases also retrieved 15 neognath protein sequences annotated as IRF3 (Suppl. Tab. 4). However, all these genes clustered in the evolutionary tree with IRF7 genes and not with IRF3 (Suppl. Fig. 2). Therefore, IRF3 genes seem to be missing in all but palaeognath birds.

### 2) Identification and sequence characterization of duck IRF9

Similar searches of genetic databases focused on avian IRF9 genes recovered multiple hits with high GC content. Such sequences were usually highly incomplete and were present on unplaced genomic contigs. In some cases (e.g. *Taeniopygia guttata* XM_041711703), NCBI automated computational analysis predicted the presence of an IRF9-like gene. Using the *T. guttata* sequence as a probe in blast searches, we detected nearly complete IRF9 in wild duck (*A. platyrhynchos*) genomic contig NOIJ01000842. Publicly available RNASeq datasets (e.g. NCBI SRA study PRJNA509092) were then used to fill the gaps and assemble a full coding sequence of duck IRF9 (Suppl. Tab. 2).

The protein sequence of the newly identified duck IRF gene was compared with sequences of other duck IRF paralogs and with complete sets of IRF genes from additional two avian, two mammalian, and one teleost species. A phylogenetic tree showed the duck gene co-localized with the other previously characterized IRF9 genes as well as with emu sequence annotated as IRF9-like in one monophyletic group, confirming its IRF9 identity (Fig. 2). The branching order of the tree validates the IRF9 genes as members of the higher order group, IRF4-G.

The multiple sequence alignment of IRF9 proteins allowed us to compare functional domains and conserved amino acid positions between different IRF9 orthologs (Fig. 3). The sequence of duck IRF9 was generally similar to protein sequences from the other species but surprisingly exhibited a high level of diversity in several regions. In a large portion of the DNA binding domain (DBD) and in several regions of the IRF association domain (IAD1), the sequences of the fish and mammalian proteins were more similar to each other than either of them is to the avian protein. Also, two of the conserved tryptophans in DBD were replaced in duck IRF9 by different amino acids, phenylalanine and cysteine. Similar replacements were described before in other IRF proteins but are relatively rare (Nehyba, Hrdlicková, and Bose 2009). Only three of the nine amino acids important for the interaction of human IRF9 with STAT2 (Rengachari et al. 2018) are conserved in duck protein.

**Figure 3.**
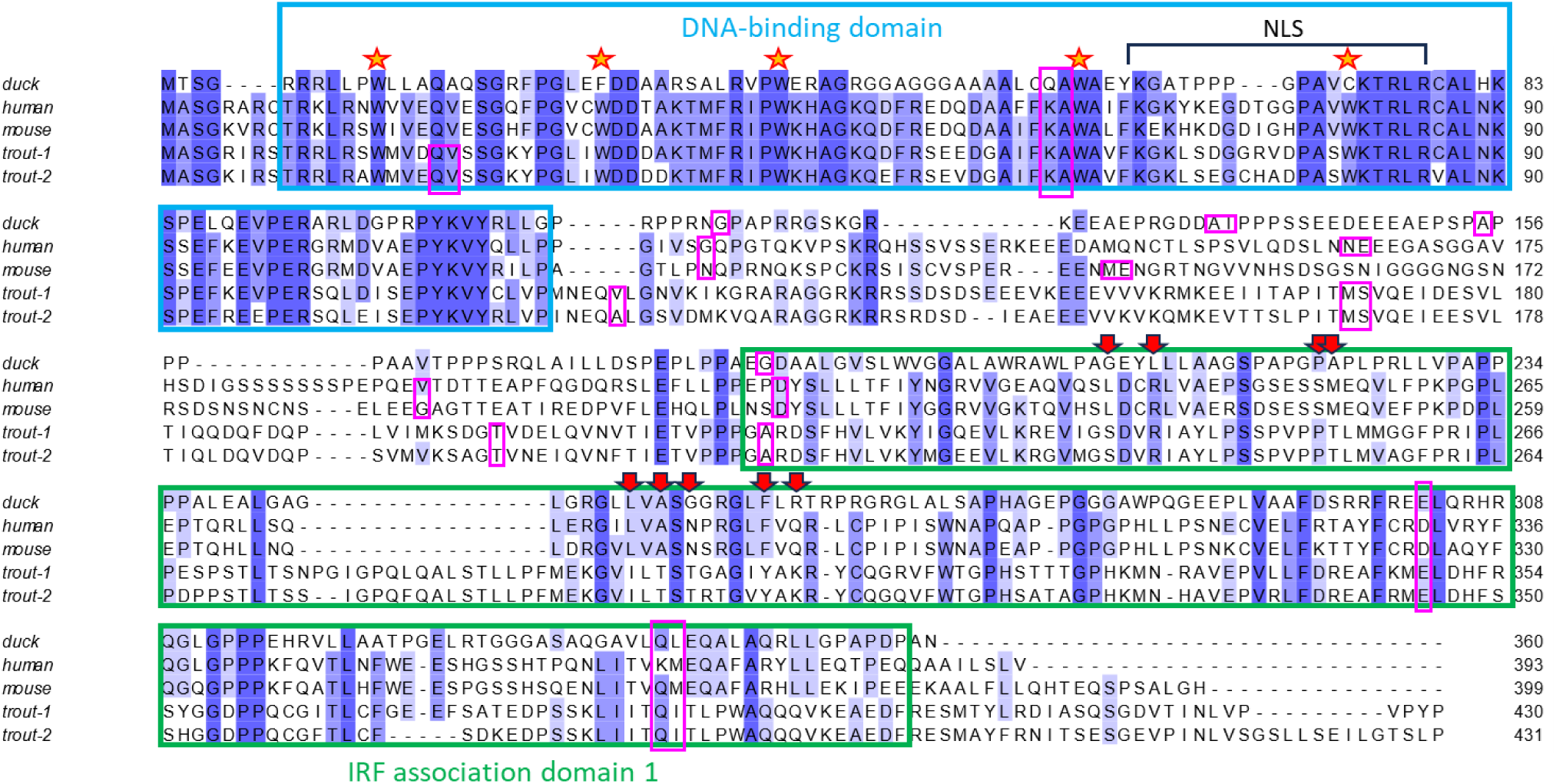
Multiple alignment of IRF9 proteins. Principal functional domains, DBD and IAD1, are designated by rectangles with blue and green outlines, respectively. The positions of five conserved tryptophans in the DBD, the prominent feature of IRF proteins, are marked by yellow stars with red boundaries. The DBD also contains a nuclear localization signal (NLS). The red arrows in the IAD1 domain designate the amino acids important for the interaction of human IRF9 with STAT2. The amino acids with codons split by introns are indicated by rectangles with purple outlines. If any intron splits CDS exactly between two codons, both of the encoded amino acids are included in the rectangle. The other sequence features are discussed in the main text. The sequences were aligned by ClustalX and the alignment visualized using Jalview 2.11.3.3. Used protein sequences are listed in Suppl. Tab. 5.

Duck IRF9 CDS is split into eight exons as is typical for other mammalian and avian genes of the IRF4-G group. Its GC content is 80%, which is in contrast with mammalian and other vertebrate IRF9 orthologs whose GC content is usually around 60%. Exon boundaries (purple outlined rectangles in Fig. 3) in the CDS are highly conserved inside DBD and IAD1, and more variable in the central linker region. The first two exons contain the sequence of the conserved DBD and short N-terminus while the last three encode IAD1 and the C-terminus of variable length. The three exons in the middle (exons 3-5) encode the DBD-IAD1 linker with low level of sequence conservation. Trout IRF9 genes have the region equivalent to the first duck/mammalian coding exon split into two exons. The introns of the duck IRF9 gene are relatively short, in total only 2.5 times longer than the CDS. Such compact gene structure is typical for most vertebrate IRF genes.

### 3) IRF9 genes are part of the gene sets of most avian species

Further iterative rounds of blast queries using the complete duck IRF9 gene sequence and multiple newly identified avian IRFs were done against NCBI databases. These searches for IRF9 genes in other avian species resulted in a substantial collection of 38 genes, including 34 full length and 4 partial CDS from 37 species (Tab. 1). These species represent 16 avian orders, including the species-rich song birds (Passeriformes), suggesting a pan-avian distribution of the IRF9 gene. Additional partial IRF9 sequences not indicated in the table were found in other song birds, but were not further analyzed. Interestingly, no IRF9 sequences were found in Galliformes despite the availability of a number of highest quality genomes from this bird order.

**Table 1.**
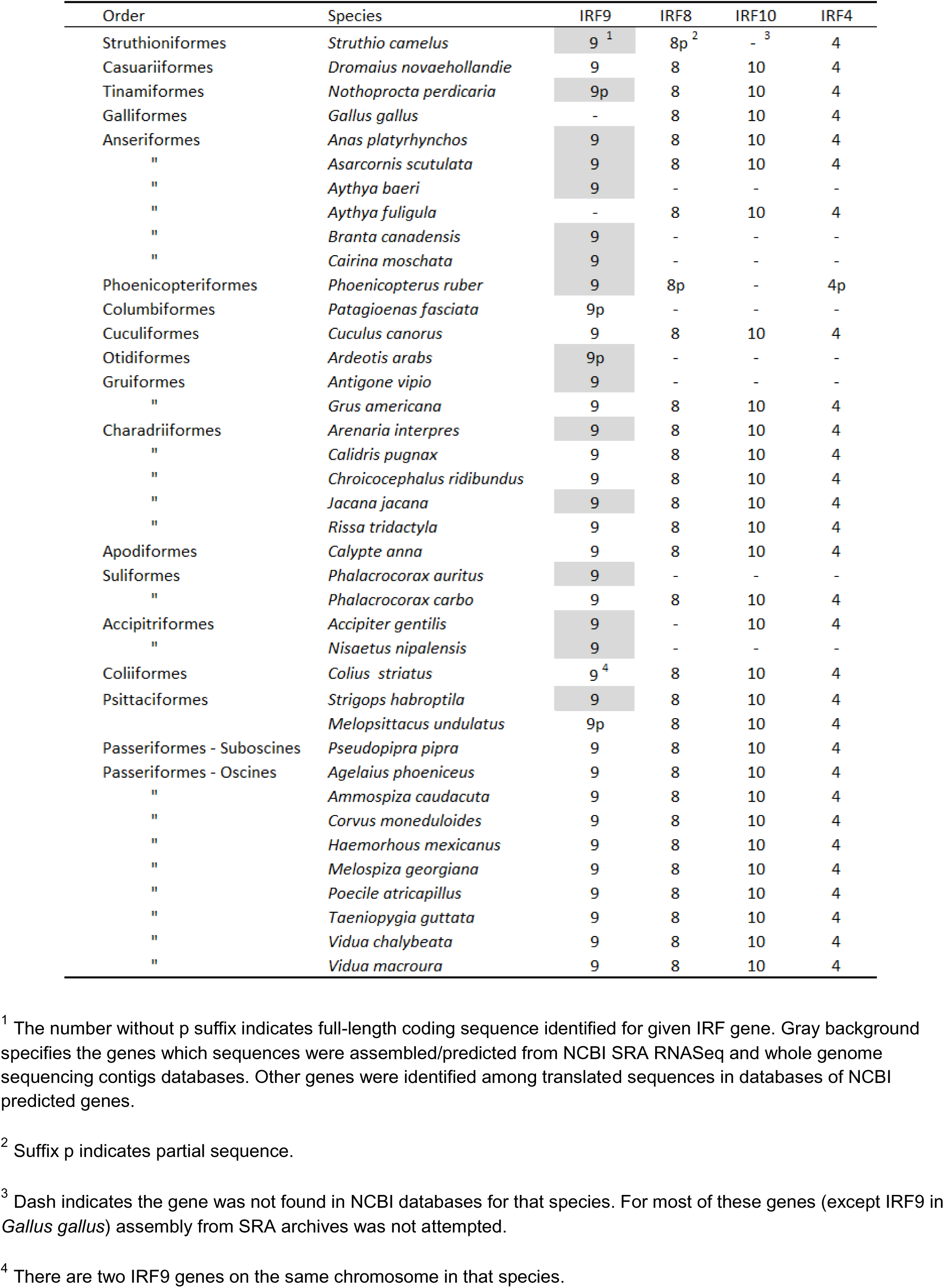
Avian IRF9 genes identified. Sequences of matching paralogs of the IRF4-G group were also retrieved in species with available annotated genomes. All sequences are included in Suppl. Tab. 5.

| Order | Species | IRF9 | IRF8 | IRF10 | IRF4 |
| --- | --- | --- | --- | --- | --- |
| Struthioniformes | *Struthio camelus* | 9 <sup>1</sup> | 8p <sup>2</sup> | - <sup>3</sup> | 4 |
| Casuariiformes | *Dromaius novaehollandie* | 9 | 8 | 10 | 4 |
| Tinamiformes | *Nothoprocta perdicaria* | 9p | 8 | 10 | 4 |
| Galliformes | *Gallus gallus* | - | 8 | 10 | 4 |
| Anseriformes | *Anas platyrhynchos* | 9 | 8 | 10 | 4 |
| " | *Asarcornis scutulata* | 9 | 8 | 10 | 4 |
| " | *Aythya baeri* | 9 | - | - | - |
| " | *Aythya fuligula* | - | 8 | 10 | 4 |
| " | *Branta canadensis* | 9 | - | - | - |
| " | *Cairina moschata* | 9 | - | - | - |
| Phoenicopteriformes | *Phoenicopterus ruber* | 9 | 8p | - | 4p |
| Columbiformes | *Patagioenas fasciata* | 9p | - | - | - |
| Cuculiformes | *Cuculus canorus* | 9 | 8 | 10 | 4 |
| Otidiformes | *Ardeotis arabs* | 9p | - | - | - |
| Gruiformes | *Antigone vipio* | 9 | - | - | - |
| " | *Grus americana* | 9 | 8 | 10 | 4 |
| Charadriiformes | *Arenaria interpres* | 9 | 8 | 10 | 4 |
| " | *Calidris pugnax* | 9 | 8 | 10 | 4 |
| " | *Chroicocephalus ridibundus* | 9 | 8 | 10 | 4 |
| " | *Jacana jacana* | 9 | 8 | 10 | 4 |
| " | *Rissa tridactyla* | 9 | 8 | 10 | 4 |
| Apodiformes | *Calypte anna* | 9 | 8 | 10 | 4 |
| Suliformes | *Phalacrocorax auritus* | 9 | - | - | - |
| " | *Phalacrocorax carbo* | 9 | 8 | 10 | 4 |
| Accipitriformes | *Accipiter gentilis* | 9 | - | 10 | 4 |
| " | *Nisaetus nipalensis* | 9 | - | - | - |
| Coliiformes | *Colius striatus* | 9 <sup>4</sup> | 8 | 10 | 4 |
| Psittaciformes | *Strigops habroptila* | 9 | 8 | 10 | 4 |
|  | *Melopsittacus undulatus* | 9p | 8 | 10 | 4 |
| Passeriformes - Suboscines | *Pseudopipra pipra* | 9 | 8 | 10 | 4 |
| Passeriformes - Oscines | *Agelaius phoeniceus* | 9 | 8 | 10 | 4 |
| " | *Ammospiza caudacuta* | 9 | 8 | 10 | 4 |
| " | *Corvus moneduloides* | 9 | 8 | 10 | 4 |
| " | *Haemorhous mexicanus* | 9 | 8 | 10 | 4 |
| " | *Melospiza georgiana* | 9 | 8 | 10 | 4 |
| " | *Poecile atricapillus* | 9 | 8 | 10 | 4 |
| " | *Taeniopygia guttata* | 9 | 8 | 10 | 4 |
| " | *Vidua chalybeata* | 9 | 8 | 10 | 4 |
| " | *Vidua macroura* | 9 | 8 | 10 | 4 |

<sup>1</sup> The number without p suffix indicates full-length coding sequence identified for given IRF gene. Gray background specifies the genes which sequences were assembled/predicted from NCBI SRA RNASeq and whole genome sequencing contigs databases. Other genes were identified among translated sequences in databases of NCBI predicted genes.
<sup>2</sup> Suffix p indicates partial sequence.
<sup>3</sup> Dash indicates the gene was not found in NCBI databases for that species. For most of these genes (except IRF9 in *Gallus gallus*) assembly from SRA archives was not attempted.
<sup>4</sup> There are two IRF9 genes on the same chromosome in that species.

The phylogenetic tree of the newly described avian IRF9 genes was constructed to establish their correct orthology (Fig. 4, Suppl. Fig. 3). Avian IRF9 protein sequences were compared with the sequences of their matching paralogs from the IRF4-G group (IRF4, 8, 10) (Tab. 1). Also, complete sets of four IRF4-G genes from several non-avian reptiles were added to the group of analyzed genes (Suppl. Tab. 5). Monophyletic clustering of the avian IRF9 sequences confirmed their correct identification. Again, chicken IRF10 gene (thick black arrow) annotated in NCBI database as IRF9 clustered with IRF10 genes and not with IRF9 genes.

**Figure 4.**
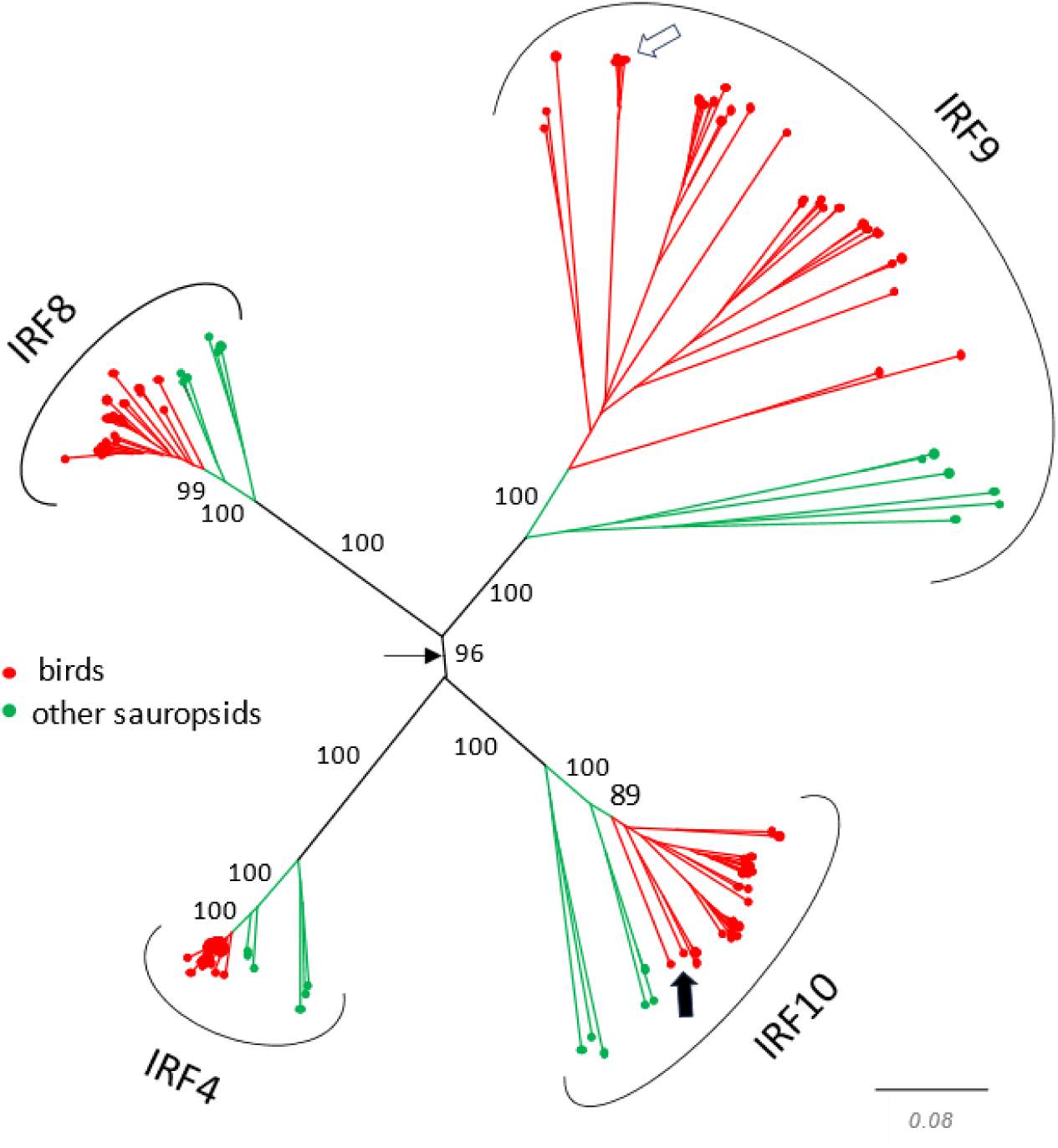
Avian IRF9 protein sequences form a single clade in the phylogenetic tree. Thirty four avian IRF9 genes with complete and two with partial CDS from those genes listed in Tab. 1 are shown in the tree. IRF4, 8, and 10 sequences from the same or closely related species (if available) and sequences from selected non-avian sauropsids are also present in the tree. The tree also contains IRF4, 8, and 10 of chicken, where IRF9 gene was not found. The sequences were either assembled/predicted from NCBI SRA or WGS archives or were identified among translated sequences in databases of NCBI assembled genes and all are listed in Tab. 1 and/or in Suppl. Tab. 5. Thin black arrow indicates the position of the root based on the hypothesis of the origin of IRF4-G genes by 2WGD. The white arrow indicates the position of anseriform IRF9 proteins that include duck IRF9. Thick black arrow shows the location of chicken IRF10 protein annotated in NCBI genomic databases as IRF9. The tree shown in this figure is also shown in the form of circular cladogram with species labels included in Suppl. Fig. 3. The NJ tree was constructed from protein sequence alignment by ClustalX. Branch support values calculated by bootstrapping (1000 replicates) are shown as percentages and only for central branches of the tree. The scale bar indicates the number of amino acid substitutions per site.

### 4) Conserved syntenic location of duck IRF9 contrasts with noncanonical chromosomal positions of IRF9 in a majority of avian species

Gene synteny in vertebrates is conservative and provides an important tool for the assessment of gene orthology. To establish if duck IRF9 is indeed co-localized on a chromosome with the same genes as in other vertebrate species, we analyzed the gene content of the longest available duck genome contig containing IRF9 (Fig. 5). All of the genes, including IRF9, found on that contig were also found on a single chromosome (chr13) in the green sea turtle (*Chelonia mydas*). In the human genome, 85% of these genes are positioned on the IRF9-containing chromosome, chr14. Duck IRF9, therefore, resides in an evolutionarily conserved chromosomal position. Similarly, at least some of those genes surrounding duck IRF9 were detected in the immediate vicinity of an IRF9 ortholog in the chromosomal sequences of *Dromaius novaehollandiae*, *Cuculus canorus*, *Grus americana*, *Phalacrocorax carbo*, *Accipiter gentilis* and *Melopsittacus undulatus*. These findings suggest that IRF9 is also located in a conserved syntenic position in six other avian orders (Casuariiformes, Cuculiformes, Gruiformes, Suliformes, Accipitriformes, Psittaciformes) in addition to the order Anseriformes. Song birds (Passeriformes), however, which represent some 60% of avian species, have the IRF9 gene located in a noncanonical position. The zebra finch (*Taeniopygia guttata*) IRF9 is on chromosome 30 while orthologs of the 13 genes that are close neighbors of duck IRF9 are instead on the zebra finch chromosome 36 (Fig. 5). The same noncanonical position is also occupied by IRF9 in other song bird species, including all of the species shown in Tab.1.

**Figure 5.**
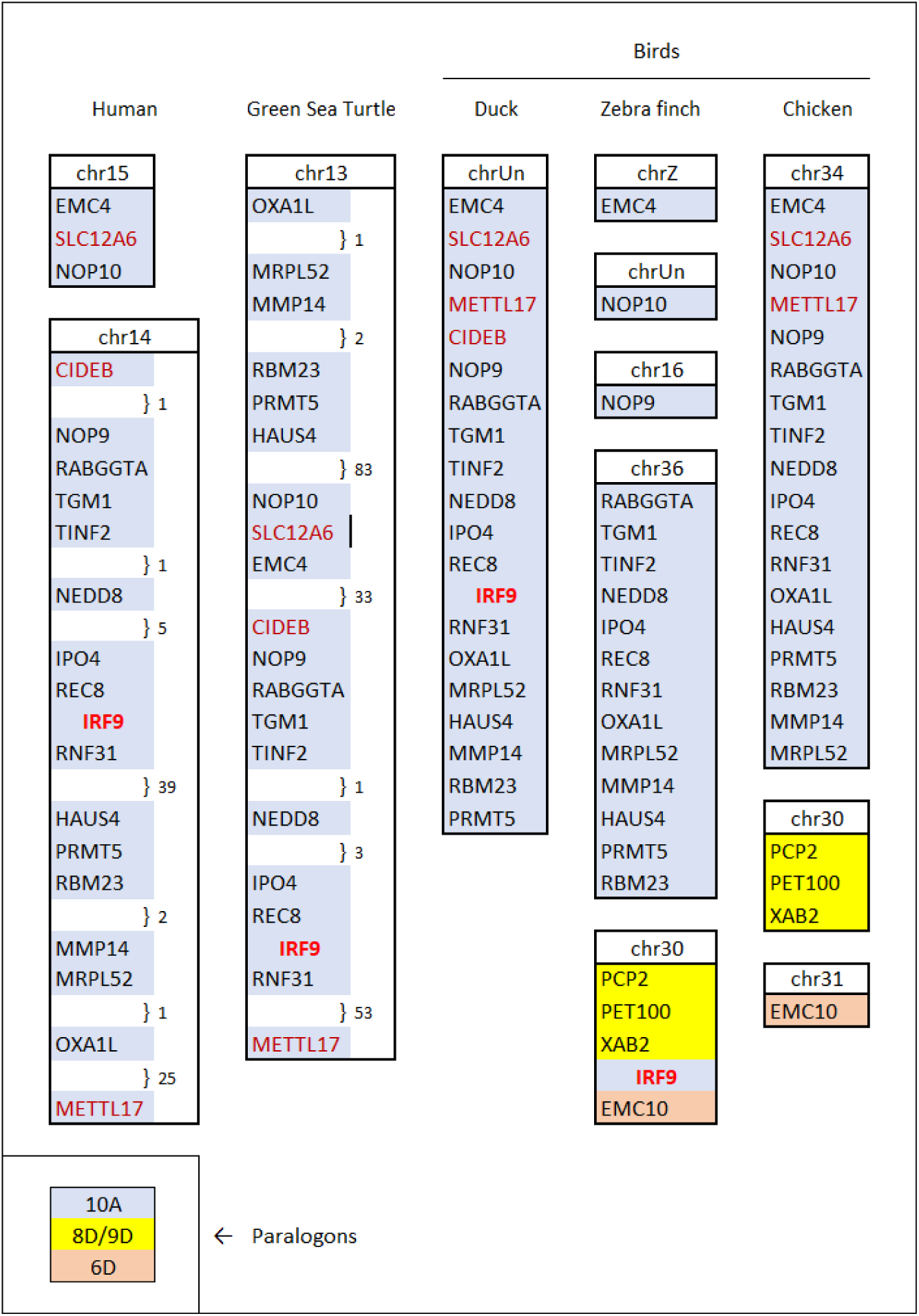
Conserved genomic position of duck IRF9, the gene relocation in song birds and the absence of IRF9 gene in galliform birds. Protein-coding genes and their positions identified in duck contig NOIJ01000842.1 on both sides of the IRF9 gene were compared with turtle and human genes (left side of the figure) and zebra finch and chicken genes (on the right). Chromosomes are numbered based on current default genomes in the NCBI database except in chicken where numbering follows the conventions of Ggswu assembly (huxu breed). The duck contig was not localized to any chromosome (chrUn). The positions of duck and chicken genes were determined by tblastn and the integrity/ completeness of their coding sequence was not verified. The same 20 genes are shown in the duck, turtle and human. Genes not found in the finch (CIDEB, METTL17, SLC12A6) or chicken (CIDEB, IRF9) are emphasized by red or purplish ink. When some protein-coding genes were omitted from the figure the gaps where they would be placed are marked by the number of genes omitted. Additional genes surrounding the relocated IRF9 are shown in finch and corresponding genes are also shown in chicken. The colored background indicates paralogon affiliation (see legend in the lower left corner). The paralogon affiliation was estimated based on the association by synteny in multiple vertebrate genomes with genes ascribed to specific paralogons by Lamb (Lamb 2021).

Gene syntenies in vertebrates are to a large extent predetermined by the two-fold whole genome duplication (WGD) that affected their ancestors in the older paleozoic (Simakov et al. 2020). Genes of jawed vertebrates can be divided into 17 groups – paralogons, each representing one chromosome of the pre-WGD chordate. Each paralogon is present in the genome in four copies designated A – D resulting in 68 gene groups identified by Lamb (Lamb 2021) and numbered from 1A to 17D. These gene groupings are surprisingly stable and still recognizable in genomes of contemporary vertebrates. The IRF9 gene is part of paralogon 10A and, by association, the other genes on the duck IRF9 contig are as well. In contrast, a majority of the genes on the zebra finch IRF9 chromosome belong to paralogons 8D/9D. Genes surrounding the zebra finch IRF9 on chromosome 30, therefore, have a completely different evolutionary syntenic origin than those of the duck. Other noncanonical positions of IRF9 were also found in five other avian orders. In each of these five cases, IRF9 is positioned in a distinct genomic context and is surrounded by genes associated with different paralogons. In Tinamiformes these are paralogons 1D/10D, in Columbiformes 5C, in Charadriiformes 3C, in Apodiformes 6C (phenomenon likely limited to hummingbirds only), and in Coliiformes 10C.

Galliform birds are the only avian order in which genes of the paralogon group 10A, including the closest IRF9-associated genes in the duck, were found but repeated attempts to find IRF9 failed. Chicken chromosome 34 where genes of group 10A reside as well as other chicken chromosomes, except W, were sequenced in their entirety from telomere to telomere (Huang et al. 2023). Nevertheless, no traces of the IRF9 sequence could be located in this or in any other chicken genome assemblies. The chromosomal locations in the chicken genome to which IRF9 was translocated in song birds (Fig. 5) and five other avian orders mentioned above are empty as well. Searches in genomes of other galliform birds also failed.

### 5) Duck IRF9 is essential factor for IFN signaling

We chose duck IRF9 for further experimental work, aiming to assess its role in the IFN signaling pathway. Previously, we generated an immortalized cell line based on duck embryonal fibroblasts (see Methods). By using CRISPR/Cas9 targeted modification in this cell line, we prepared a duck IRF9 knockout cellular clone (KO) (Fig. 6) and compared its responsiveness to duck IFN-α with a wild type (WT) clone. As a readout of IFN signaling, we measured the mRNA induction of two duck ISGs: interferon induced protein with tetratricopeptide repeats 5 (IFIT5) and myxovirus resistance protein (Mx). For these experiments, we also prepared duck IRF9 (dIRF9) expression plasmid with N-terminal flag tag. As this construct was prepared by commercial gene synthesis, the codons were optimized to avoid GC-rich stretches in IRF9 sequence, but to fully conserve the protein sequence (Suppl. Fig. 4).

**Figure 6.**
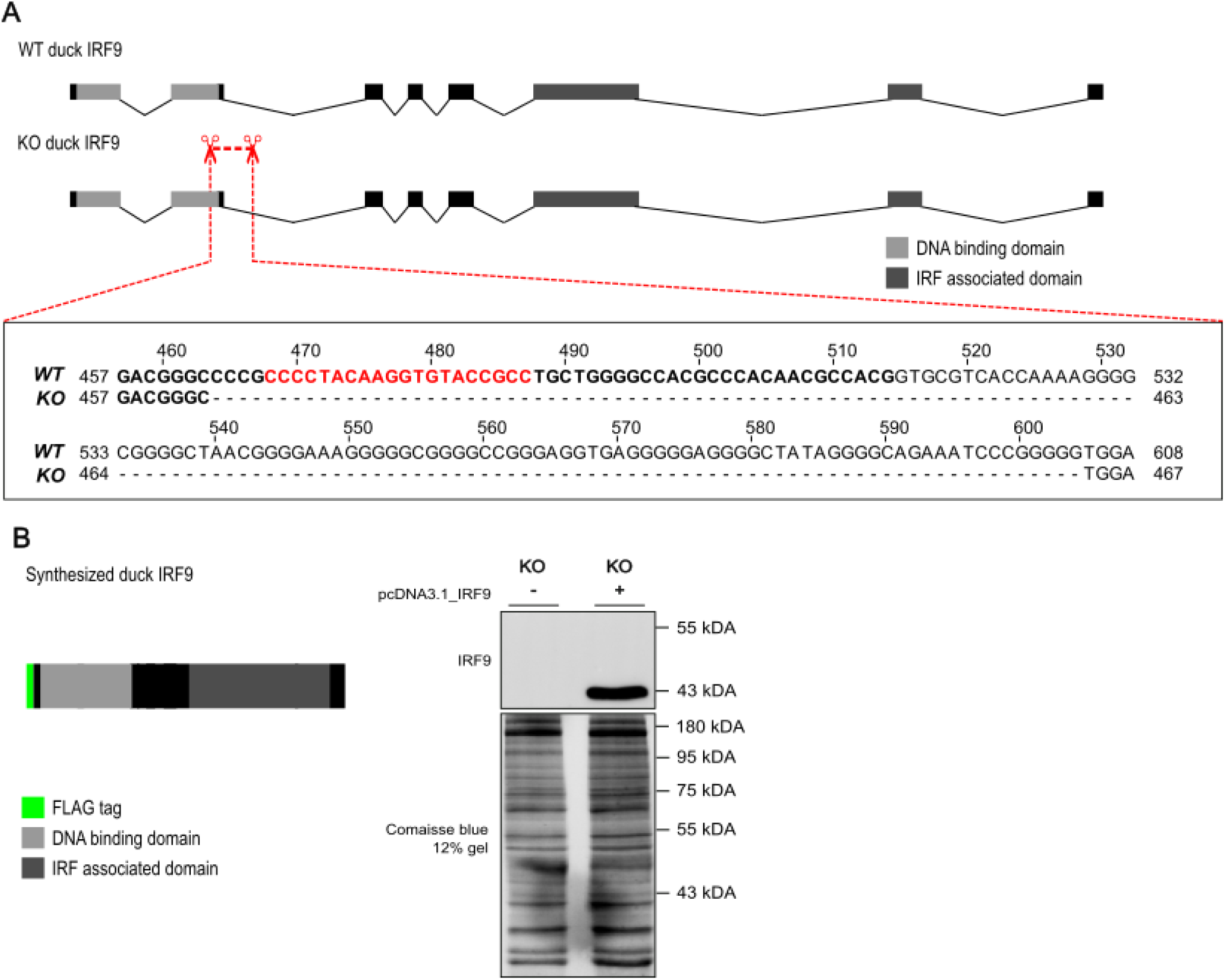
Duck IRF9 knockout and in vitro synthesized construct. A) Schematic depiction of duck IRF9 exon structure in wild type (WT) and knockout (KO) cells. Position of the deletion in the KO clone is highlighted and expanded in sequence below. Bold text represents the exon sequence and regular font the intronic part of the gene. Red font represents the position of gRNA used to target the CRISPR/Cas9 complex. Numbering of the alignment starts at the start codon of the CDS and includes introns. B) Scheme of the protein encoded by the in vitro synthesized duck IRF9 tagged with FLAG tag at N-terminus; western blot of lysate from KO cells transiently transfected by the dIRF9 expressing plasmid detected by anti-FLAG antibody.

Duck IFN caused high induction of both IFIT5 and Mx mRNA expression in WT cells (Fig. 7). This induction was strongly decreased in duck IRF9 KO cells. Further, transfection of the dIRF9 expression plasmid into KO cells partially restored the ISG expression. The partial restoration of IFN sensitivity is consistent with the fact that IRF9 was delivered by transient transfection, and therefore not all cells were targeted. Overall, these results confirm that duck IRF9 is essential for the IFN signaling pathway.

**Figure 7.**
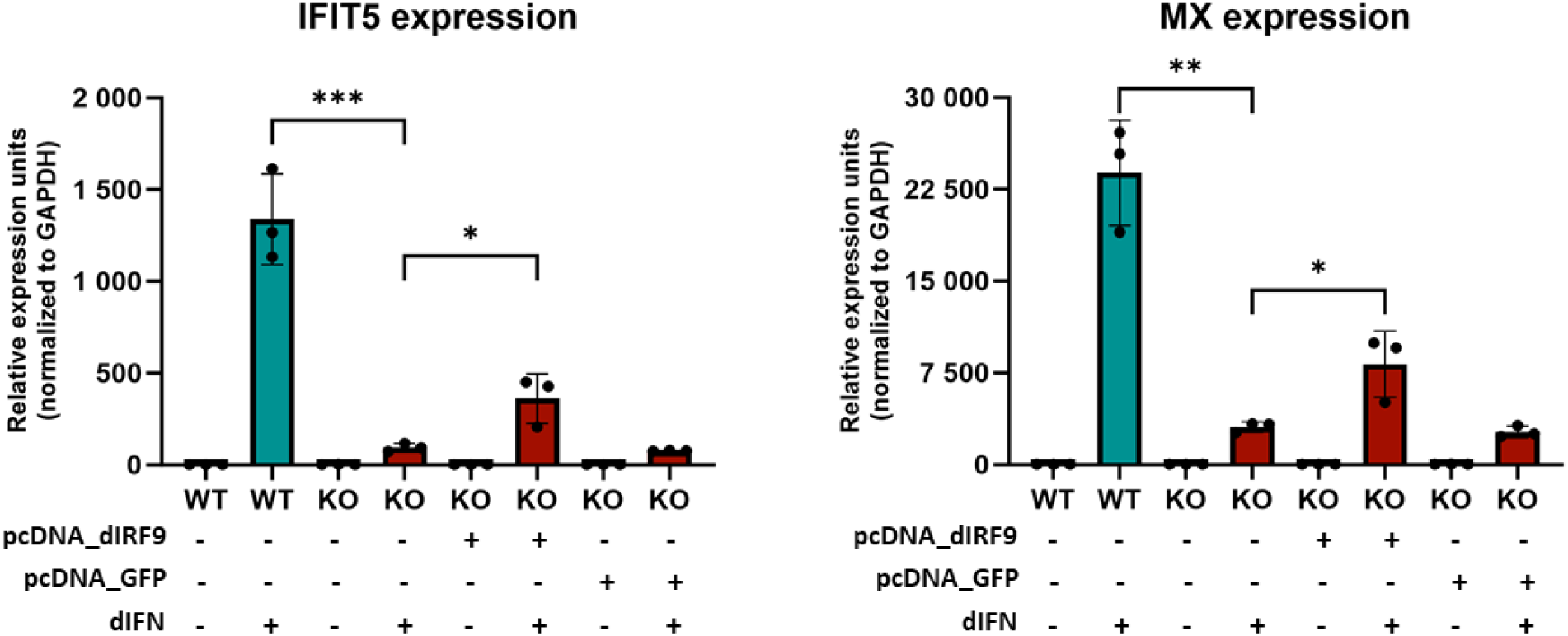
Expression of ISGs after IFN stimulation in IRF9 knockout cells. WT and KO cells were either transfected with plasmid containing in vitro synthesized duck IRF9 (dIRF9) or with green fluorescent protein (GFP)-expressing plasmid used as a negative transfection control, or they remained untransfected. Relative mRNA expression (normalized to GAPDH expression) of interferon stimulated genes IFIT5 and Mx after duck IFN (dIFN) treatment was measured by quantitative PCR. Graphs and nested t-tests were done in Graphpad Prism 10. * p ≤ 0.05; ** p ≤ 0.01; *** p ≤ 0.001.

### 6) Both IRF9 and intact ISRE are required for high ISG induction

Next, we asked whether the ISG induction pathway in duck cells requires intact ISRE. For this purpose, we used a luciferase reporter construct preceded by the promoter region of duck tetherin gene. Tetherin (also called bone marrow stromal antigen 2; Bst2) is a highly IFN-induced avian gene, which we previously characterized in chicken and other avian species (Krchlíková et al. 2020, 2023). We generated two variants of the tetherin-luciferase construct: i) with native ISRE sequence (complying with the ISRE consensus; (Santhakumar et al. 2018)) and ii) a mutant with a key guanine changed to adenine (Fig. 8). Upon transfection into WT duck cells followed by dIFN addition, the construct with intact ISRE drove high luciferase reporter expression. Only low luciferase activity was detected in the construct with mutant

**Figure 8.**
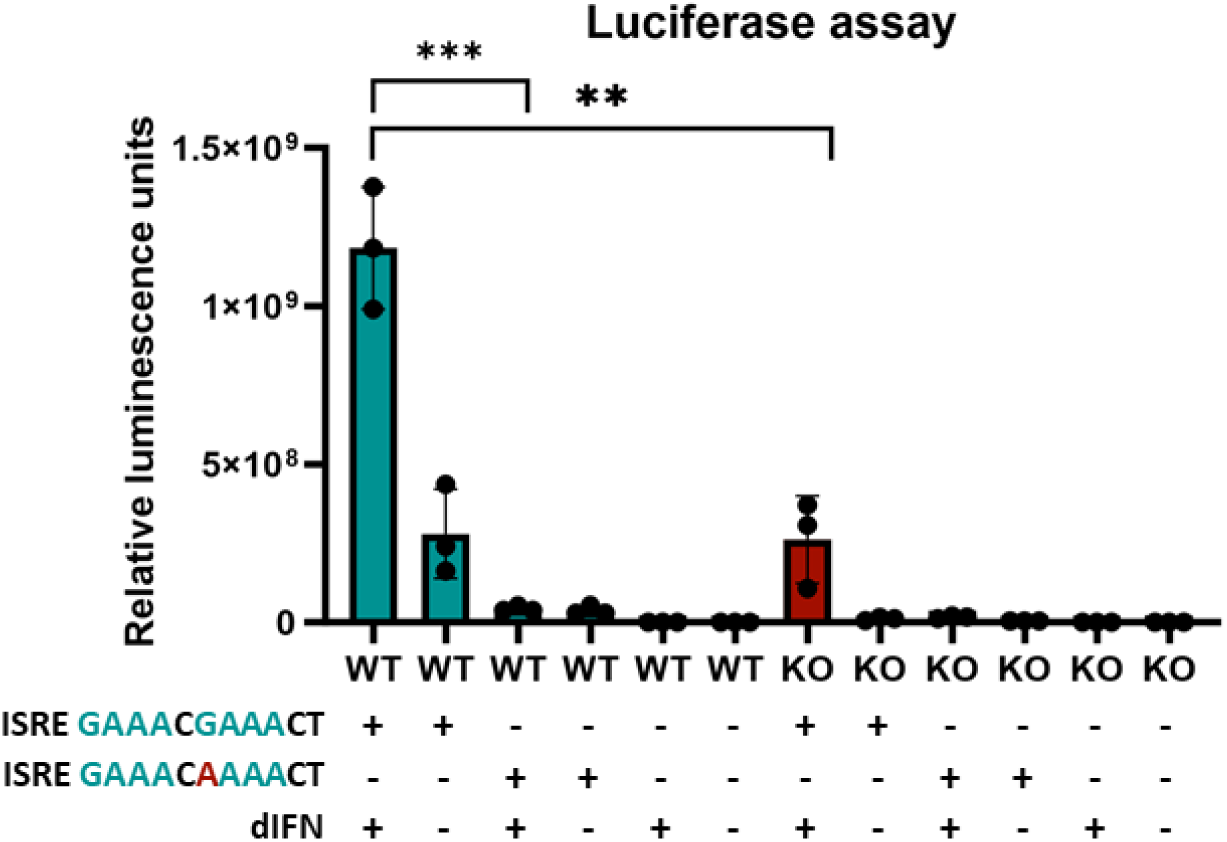
Duck IRF9 and consensus ISRE sequence is needed for expression of ISGs. Duck IRF9 WT and KO cells were transfected with pNL1.2NLucP plasmids containing ISRE–dependent tetherin promoter region and NanoLuc luciferase reporter gene. One plasmid had a promoter region containing consensus ISRE sequence GAAACGAAACT. Second plasmid had a promoter region containing the ISRE sequence with a point mutation G>A; GAAACAAAACT (nucleotide substitution highlighted in red font). Upon transfection with ISRE constructs and dIFN stimulation for 24h, the luciferase activity was determined. Graphs and nested t-test were done in Graphpad Prism 10. ** p ≤ 0.01; *** p ≤ 0.001.

ISRE. Consistent with previous experiments, expression of the ISRE-driven reporter was lower in IRF9 KO cells. These experiments show that ISG expression in duck is dependent on intact ISRE and that both intact ISRE and functional IRF9 are required for full IFN-induced ISG induction.

### 7) IRF9 is necessary for IFN-dependent protection against VSV cytopathic effect

The importance of duck IRF9 in the IFN signaling pathway culminating with ISG induction was established above. We also assessed the role of IRF9 in protection against virus infection. To achieve this, we used a well-established model of vesicular stomatitis virus (VSV)-induced cytopathic effect. The cytopathic effect can be prevented by IFN treatment in the context of a functional IFN signaling pathway (Marcus et al. 1977). This has also been used as a basis for determining IFN activity (Pestka et al. 1987). We compared the ability of IRF9 KO and WT cells to be protected against VSV-mediated cell killing by duck IFN. As a positive control, we used a supernatant from duck splenocytes stimulated with R848 (a toll-like receptor ligand), which induces multiple cytokines including interferons. Indeed, while the WT cells could be protected by IFN addition against the VSV cytopathicity, the IRF9 KO cells showed significantly lower survival (Fig. 9).

**Figure 9.**
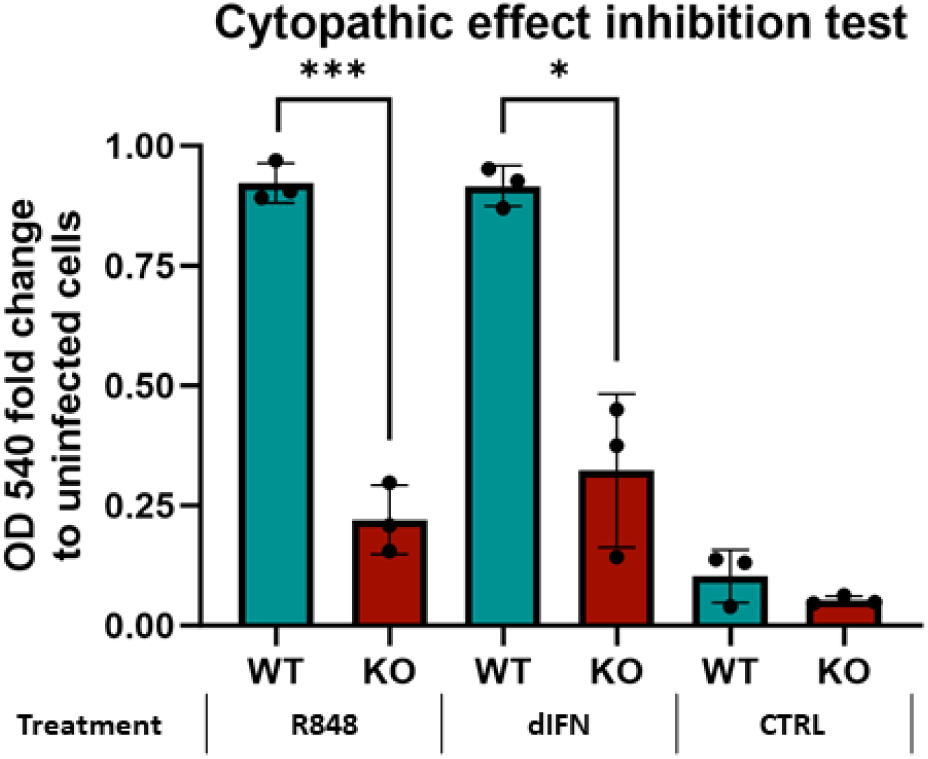
Role of IRF9 in protection of duck cells against the cytopathic effect of VSV after IFN stimulation. Duck IRF9 WT and KO cells were incubated with the following: a medium with supernatant from R848-stimulated duck spleen cells (positive control) or duck interferon supernatant (dIFN) produced in HEK293T cells or with control supernatant (CTRL). After 24h cells were infected with VSV and 24 hpi and after addition of neutral red for 2h, OD was measured using ELISA reader at a wavelength 540 nm indicating cell survival. WT cells showed significantly higher survival after incubation with R848 and dIFN compared to KO cells. Graph and Welch’s t-test were done in Graphpad Prism 10. * p ≤ 0.015; ** p ≤ 0.01; *** p ≤ 0.001

## DISCUSSION

In this work we identified two avian interferon regulatory factors, IRF3 and IRF9. Both were not characterized before and were considered missing in chicken and by extension in all birds. All the newly identified IRF3 and IRF9 orthologs are GC-rich. Therefore, their delayed identification is in line with the previous cases of “hidden genes” described by us and by others (Hron et al. 2015; Huttener et al. 2021). Despite the new identifications, our searches showed that the orthologs of the two IRFs are not present in all avian species. IRF3 orthologs are more narrowly distributed and were found only in the basal avian infraclass Palaeognathae, which includes ostriches, kiwis etc. In contrast, IRF9 orthologs are present in almost all avian species with the notable exception of the order Galliformes, which includes the domestic chicken.

It remains formally possible that even in the instances where we currently predict an IRF gene loss, the respective IRF ortholog could still be in fact present (e.g. IRF9 in galliform birds or IRF3 in Neognathae). This could happen because the extremely difficult DNA sequence of these genes (high GC-content, complex secondary structure) cannot be processed by any current sequencing technology. Alternatively, the sequence divergence of the respective IRF ortholog could be so large that it would be missed by our homology-based searches. Indeed, avian IRF9 gene sequences diverged more extensively than sequences of any other IRF. Nevertheless, given the largely unbiased nature of new long-read sequencing platforms, especially Oxford Nanopore (Browne et al. 2020), and the sensitivity of blast-based searches even for distant relatives (Suppl. Tab. 3), we consider the existence of such hidden IRFs rather improbable.

IRF3 identified only in the infraclass of Palaeognathae was always located on the chromosome next to SCAF1, in the syntenic position conserved at least from the last common ancestor of all contemporary gnathostomian vertebrates. In neognath species that gene position beside SCAF1 was found empty. Similarly, IRF9 genes in anseriform duck and in representative species of many other avian orders were located in evolutionary conserved position, i.e. between RNF31 and REC8. In galliform birds, however, no traces of IRF9 were found between these two genes nor in any other place in the genome. Surprisingly, in various species of about half of the avian orders studied, the IRF9 was found located in several noncanonical chromosomal positions indicating past translocations in the course of avian evolution. These translocations were not the relatively common intrachromosomal gene reshufflings but transfers to a different chromosomal context indicated by the repositioning into unrelated genetic linkage groups of ohnologs. Phylogenetic analysis confirmed that all these translocated genes are true IRF9 orthologs and not more distantly related members of the IRF family (Fig. 4). To our knowledge such level of synteny variability is unprecedented for vertebrate genes. While the evolutionary mechanisms of IRF3 and IRF9 gene losses and translocations are not known, we suspect that these phenomena are related. They could be the consequence of these genes being located before the loss/translocation on the smallest avian chromosomes designated as dot chromosomes. Dot chromosomes have a very peculiar chromatin structure and suffer the highest rate of recombination (Huang et al. 2023; Groenen et al. 2009). Mechanisms of the IRF3/IRF9 gene losses also seem different from the described loss of IRF10 by pseudogenization in many mammalian species as we found no traces of any avian IRF3 or IRF9 pseudogenes (Li et al. 2023).

IRF genes are a typical example of the gene family that expanded due to two whole genome duplications (2WGD) early in the vertebrate evolution. While high retention rate of IRF paralogs after 2WGD suggests that the paralogs diversified their functions, a certain level of redundancy in function among different paralogs still could be expected. Accordingly, IRF3 function in the IFN induction pathways largely overlaps with IRF7 (Servant, Tenoever, and Lin 2002; Honda, Takaoka, and Taniguchi 2006; Sato et al. 1998). Therefore, it has been suggested that in chicken, the role of IRF3/7 was substituted solely by IRF7 (Cheng et al. 2019; Guabiraba et al. 2024). As in paleognath birds both IRF3 and IRF7 are present, it will be interesting to dissect their roles experimentally in the future (Fig. 10). For example, is ostrich IRF3 expressed rather constitutively and IRF7 is IFN-inducible as they are in mammals?

**Figure 10.**
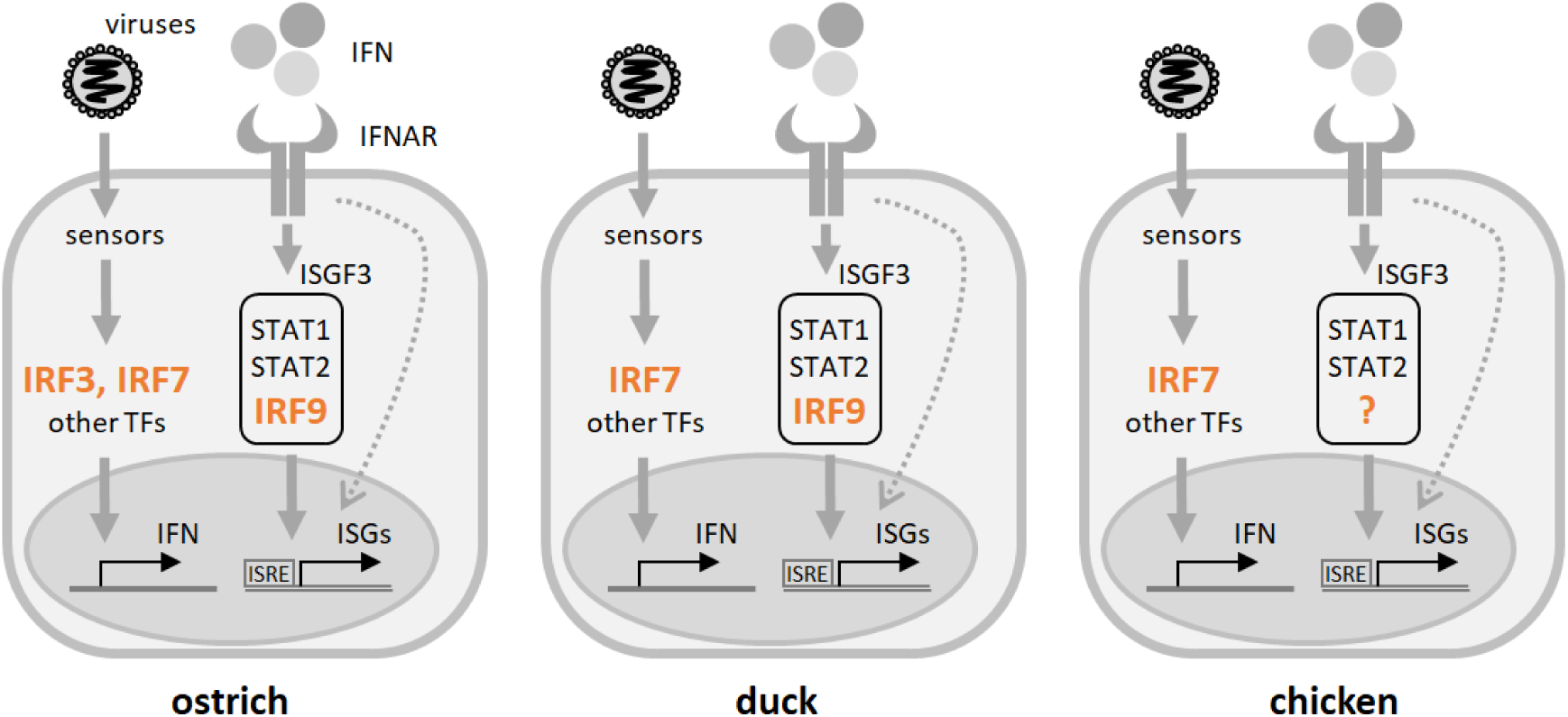
IFN induction and signaling in three avian species. Simplified schematic depiction with emphasis on the role and presence of IRF3, IRF7 and IRF9. In ostrich (*S. camelus*), all three genes are present, representing the situation in paleognath birds. Chicken (*G. gallus*) and other galliform birds have only IRF7. Duck (*A. platyrhynchos*) lacks IRF3 and serves as a representative example of a majority of birds except palaeognaths and galliforms. The dotted arrow shows various ISGF3-independent modes of ISG induction. TF – transcription factors; IFNAR – type I IFN receptor; for other abbreviations see text.

IRF9 has a unique role in the ISGF3 complex with no known substitutes in vertebrates. Therefore, its absence in chicken and other galliform birds is quite intriguing (Fig. 10). As mentioned above, it remains possible that an extremely diverged/difficult sequence of chicken IRF9 translocated into noncanonical syntenic position exists. In our opinion, it is much more probable that another member of the chicken IRF family can be involved in the ISGF3 complex formation. Alternatively, chicken IFN signaling might involve alternatives to classical ISGF3. Many IFN signal transduction modes independent of classical ISGF3 complex formation have been described (Platanitis et al. 2019; Majoros et al. 2017).

To elucidate if the avian IRF9 still retains functions in the IFN system described for this gene in mammals we performed a series of experiments. For this work, we selected duck IRF9, because the order Anseriformes that includes duck species represents the closest relationship to galliform birds. Knockout experiments in duck embryonal fibroblasts showed that, at least in this cell type, IRF9 is essential for IFN-mediated induction of ISGs. Interestingly, residual induction of ISGs is occurring even in knockout cells. This contrasts with mammalian IRF9 knockouts, where ISGF3-dependent ISG expression was mostly negligible (Skavicus and Heaton 2023). For now we can only hypothesize that the IRF9-independent induction suggests the existence of ISGF3 alternatives, possibly relevant also for the situation in IRF9-deficient chicken cells. As expected, a canonical ISRE sequence is required, together with intact IRF9, for full induction of ISGs in duck cells (Fig. 8). We suppose that duck ISGF3 binds to ISRE through IRF9-DNA interactions, similarly to mammalian ISGF3 (Darnell, Kerr, and Stark 1994; Nowicka et al. 2023). Finally, we proved the importance of duck IRF9 in a broader context than the ISG induction pathway. For this, we used the protective effect of IFN on VSV cytopathicity, a classical assay used to titer IFN activity (Fig. 9). Intact IRF9 was shown to be essential for IFN-induced antiviral responses in duck cells. This is in agreement with results obtained using mouse IRF9 knockouts (Kimura et al. 1996). In summary, we have identified two previously uncharacterized members of the avian IRF family with key roles in the IFN signaling pathways. Further research will show the details of their function and the adaptations present in avian species lacking specific IRF orthologs.

## METHODS

### Computational analysis of IRF genes in avian species

Sequences in the NCBI database (whole genome shotgun (WGS) genomic assemblies, nonredundant (nr) database, SRA sequences) were searched with various IRF sequences using BLAST (Camacho et al. 2009). The sequences obtained were downloaded and assembled manually either with CLC genomics workbench 21.0.5 (Qiagen) or with Lasergene 10.0.0 (DNASTAR, Madison, WI, USA). Partial CDSs from avian genomic contigs were, if possible, completed using SRA NCBI data.

### Cell lines and culture conditions

HEK293T cell line was cultured in 5% CO_2_ atmosphere at 37°C in medium containing Dulbecco’s modified Eagle’s medium (DMEM, Sigma-Aldrich) supplemented with 7 % fetal calf serum (Gibco) and 100 μg/ml penicillin and streptomycin. Duck fibroblast cell lines were cultured in 5% CO_2_ atmosphere at 37°C in medium containing mixture of 1:1 (DMEM)/F-12 (Sigma-Aldrich) supplemented with 4% calf serum (Gibco), 4% fetal calf serum (Gibco), 1% chicken serum (Gibco) and 100 μg/ml penicillin and streptomycin.

Immortalized duck embryo fibroblasts (DEF) were prepared by standard procedure from 14th-day old embryos of inbred Khaki Campbell ducks (Stepanets et al. 2001). For propagation of cells, DMEM/F12 (Gibco) medium supplemented with 5% of calf serum, 5% of fetal calf serum, 1% of chicken serum, 1% of vitamins (Sigma), 1% non-essential amino acids (Sigma), 1% folic acid (Sigma), 1% Glutamax 100 (Gibco) and 0,5% dimethyl sulfoxide was used. Secondary DEF were transfected with pRNS1 plasmid (Litzkas, Jha, and Ozer 1984) containing simian virus 40 (SV-40) DNA sequences by Lipofectamine 2000 reagent (Invitrogen) according to the manufacturer’s instructions. Two days later transfected cells were treated with a fresh medium containing G418 selection antibiotics (Sigma). A week later the first colonies of neomycin resistant cells appeared in transfected culture. Starting the 14th day after transfection cells were regularly passaged twice a week up to the passage 30. Then followed a period of slowed growth, cells became highly senescent, granulated and vacuolated with a cell detritus in the culture medium. In this period the cells had to be transferred at high density, but since the passage 36 cells started to proliferate again and were regularly passaged three times a week. At the passage 98 a stock of cells fully established for growth in vitro was frozen. Since the cells exhibited very heterogeneous morphology two rounds of cloning were performed with the aim to get cell population resembling secondary duck fibroblasts. In this way a clone D7 was obtained that was further used in knockout experiments.

### Generation of the duck IRF9 knockout cell line

To create an IRF9 knockout duck cell line we used standard CRISPR/Cas9 technique as described before on avian cells (Koslová et al. 2018). Duck cells (above) were seeded at a density of 8 x 10^5^ on 60 mm wells and grown to approximately 80% confluence. The gRNA-encoding sequenced were prepared as sense and antisense oligonucleotides (5’-CACCGGCGGTACACCTTGTAGGGG, 5’-AAACCCCCTACAAGGTGTACCGCC) specific to sequence at the end of duck IRF9 exon 2, mixed, phosphorylated, denatured, annealed and ligated into pX458 plasmid (Ran et al. 2013). The resulting plasmid was transfected using Lipofectamine 3000 (Invitrogen). Two days after transfection, single cell sorting into 96 well plates using Influx cell sorter (Becton, Dickinson) was done to create GFP-positive clonal cell lines. Viable clones were expanded to sufficient size for standard phenol-chloroform protocol DNA isolation. These clones were genotyped for successful homozygous Cas9 cleavage in targeted region using PCR (primers 5’-GGCGTGGGCGGAGTACAAAGG and 5’-GGCTCGGCTCCGCCTCC); PCR products were verified by Sanger sequencing. For further experiments we chose a clone with large homozygous 141 nucleotide deletion over a splice site that deletes 56 nucleotides in the exon region. As negative control we chose a clone where no mutation occurred.

### Cloning and expression of duck IFN-**α**

Duck IFN-α CDS (accession number X84764.1), an intronless gene, was amplified from genomic DNA with primers containing restriction sites *BamH*I and *EcoR*I. The digested PCR product was then cloned into a pGEM-T Easy plasmid (Promega) and verified by Sanger sequencing. To generate the final expression construct, the insert was re-cloned into a pcDNA-3.1 plasmid (GenScript Biotech Corporation). HEK293T cells were seeded into 10-cm plates overnight to almost full confluence. They were then transfected using standard Lipofectamine 3000 protocol (Invitrogen) with pcDNA 3.1 plasmid containing duck IFN-α or using the same plasmid with GFP insert instead as a negative control. 24 and 48 hrs post transfection, medium was collected and pooled resulting in one stock of supernatant containing duck IFN and one stock of control supernatant. The supernatants were cleared by centrifugation and filtering (0,45 μm) and stored at –20°C.

### Analyzing the role of duck IRF9 in expression of ISGs

Duck WT and IRF9 KO cells were seeded at a density of 5 x 10^5^ cells per well of 6 well plates and grown to approximately 80% confluence. Cells were treated as following: KO cells were transfected using standard Lipofectamine 3000 protocol (Invitrogen™) either with pcDNA 3.1 plasmid containing duck IRF9 fused with FLAG tag on N-terminus (GeneScript) or control pcDNA 3.1 plasmid containing GFP or remained without transfection. Approximately 18 hrs post transfection, either supernatant containing duck IFN was added in a final dilution 1:1000 (v/v) to both WT and KO cells or the cells remained untreated. 6 hrs after IFN treatment cells were collected for RNA isolation. Experiment was replicated twice independently; each experiment was done in three biological replicates for each experimental condition. Ectopic expression of tagged dIRF9 was verified by western blot using mouse anti-FLAG antibody (Sigma).

### RNA isolation, reverse transcription and quantitative PCR

Total RNA was isolated from cultured cells using RNAzol RT (Molecular Research Center) according to the manufacturer’s protocol. Reverse transcription was then performed with 0,5-1 μg of RNA using ProtoScript II First Strand cDNA Synthesis Kit (NEB) and 3′-RACE CDS primer (Clontech). Next, quantitative PCR was done using MESA GREEN qPCR MasterMix Plus (Eurogentec). Primers targeting duck IFIT5 (5’-AAGCTACCTTCAAACGGGTA and 5’-TCCTCCTTCAGCAAAGTCCA), Mx (5’-TCATGACTTCGGCGACAAC and 5’-AACTCGGCCACTGAGGTAAT) and GAPDH (glyceraldehyde-3-phosphate dehydrogenase; 5’-TGTCTCCTGCGACTTCA and 5’-TCCTTGGATGCCATGTGGAC) were used for expression analysis. Specificity of PCR products was verified by melting curve analysis. Each sample was analyzed in technical triplicates.

### Luciferase assays

The duck tetherin promoter region (accession number VSDN01000029.1; range:1112681-1112965) was amplified from genomic DNA. The PCR product was then digested with primer-derived *Kpn*I and *Xho*I restriction enzymes and inserted into the *Kpn*I/*Xho*I-digested pNL1.2-NLucP vector (Promega). The substitution G>A into the ISRE consensus sequence was introduced by In-Fusion cloning (TaKaRa). The promoter region was amplified in two fragments. The fragments were then cloned together into the pNL1.2NLucP plasmid using In-Fusion Cloning Kit according to the manufacturer’s protocol. Duck WT and IRF9 KO cells were seeded at a density of 5 x 10^5^ per well into 6 well plates and grown to approximately 80% confluence. The cells were then transfected with plasmid containing NanoLuc luciferase reporter gene and a promoter sequence either with consensus ISRE (GAAACGAAACT) or with mutant ISRE (GAAACAAAACT). 24 hrs post transfection cells were stimulated with dIFN supernatant with final dilution 1:1000 (v/v) for another 24 hrs. For luciferase signal measurement, Nano-Glo® Luciferase Assay System (Promega) mix of lysis buffer and substrate was added to 1 x 10^5^ cells. Lysate was then transferred to white plates for luminescence readout. The experiment was performed in biological triplicates and each replicate was measured in technical triplicates.

### Cytopathic effect inhibition assay

Cytopathic inhibition effect of duck WT and IRF9 KO cells following viral infection was tested according to Pestka et al. (Pestka et al. 1987) with some modifications. The cells were seeded at a density of 1 x 10^4^ cells/100 µl into 96-well plates and grown in the presence of the following additives: medium with supernatant from R848 (InvioGen)-stimulated duck spleen cells as a positive control (diluted 1:20 v/v), medium with supernatant containing duck IFN (see above, diluted 1:2000 v/v) and medium with control supernatant as a negative control. After 24 hrs, cells were infected with Vesicular Stomatitis Virus (a gift from Mathias Büttner, Ludwig Maximilian University) with final dilution 1:2000 and incubated for 24 hrs until a clear cytopathic effect was seen in the control cells with control medium only. All media were removed from the cells and replaced by fresh medium containing neutral red at a concentration of 0,01%. After 2 hrs cell culture supernatants were removed and cells were gently washed, dried and 50 µl of a 3M guanidine hydrochloride solution was added to solve the pinocytosed neutral red. OD was measured using an ELISA reader at a wavelength of 540 nm. Three independent assays were performed; results represent the mean of the three assays.

### Statistics

For qPCR and luciferase assay results, we used nested t-test and for cytopathic effect inhibition assay we used Welch’s t-test, both implemented in Graphpad Prism.

## ACKNOWLEDGEMENTS

This work was funded by grant GA23-07210S (to D.E.) from the Czech Science Foundation. D.E., J.H., J.N., D.K. and V.K. were further supported by the project National Institute of virology and bacteriology (Programme EXCELES, No. LX22NPO5103) funded by the European Union—Next Generation EU. We also acknowledge institutional support from project RVO 68378050. This project was partially funded by the Deutsche Forschungsgemeinschaft (DFG, German Research Foundation) in the framework of the Research Unit ImmunoChick (FOR5130) project KA 1936/3-1 (to B.K.).

