## Supplementary figures for "Avian interferon regulatory factor (IRF) family reunion: IRF3 and IRF9 found"

```
dnoIRF3         MGAPKPLILPWLRAQLDRGGYPGVRWLG--ATRFRVPWKHGLRHDVAPEDFQLFQDWAIASGCYRPGVDPPAPAVWKRNFRAALNRKPGLRVVEDRSGDAGDPHKVYEFLPGGPGGAGDP
scaIRF3         MGTPKPLILPWLRAQLDRGCYPGVRWLG--ATRFRVPWKHGLRHDVAPEDFQLFQDWAIASGCYRPGVDPPVPAVWKRNFRSALNRKPGLRMLEDHSGDANDPHKVYEFLA---------
rpeIRF3         MGNPKPLILPWLRAQLDRGTYPGVRWLG--GTRFRVPWKHGLRHDVAPEDFQLFQDWAIASGCYRPGVDPPAPAVWKRNFRAALNRKPELRVVEDRSGDAGDPHKVYEFLP---------
ccarIRF3        MGTPKPLIVPWLRKKLDDGCYPGVSWLDQGRTQFRVPWKHGLRQDASSEDFQLFRDWAIDSGNYRPGHDAPTPSIWKRNFRSALNRKPGIQVLQDHSSDSADPHKVYEILP---------
asiIRF3         MGSQRPLLVPWLREQLDSGCYPGVCWLNAERTRFRVPWKHGLRHDAAREDFQLFQDWATVSGCYHAG-TAPVPSIWKRNFRSALNRKPEFKVVEDNSSDATDPHKVYEILP---------
hsaIRF3         MGTPKPRILPWLVSQLDLGQLEGVAWVNKSRTRFRIPWKHGLRQDAQQEDFGIFQAWAEATGAYVPGRDKPDLPTWKRNFRSALNRKEGLRLAEDRSKDPHDPHKIYEFVN---------
cfaIRF3         MGTPKPRILPWLVSQLDLGQLEGVAWLDERRTRFRIPWKHGLRQDAQQEDFGIFQAWAEVSGAYTPGKDKPDLPTWKRNFRSALNRKEELRVAEDRSKDPHDPHKVYEFVI---------
mmuIRF3         METPKPRILPWLVSQLDLGQLEGVAWLDESRTRFRIPWKHGLRQDAQMADFGIFQAWAEASGAYTPGKDKPDVSTWKRNFRSALNRKEVLRLAADNSKDPYDPHKVYEFVT---------
                *   :* ::***   :** *    ** *:.    *:**:*******:*.    ** :*: **  :* * .*    *   . ******:*****  :::   *.* *.  ****:**::

dnoIRF3         PEVYEFLPGALGGASGAGDPPETYEFLPGVPGGASGAGDPPEVYEFLPGVPGGAGQPPRAGDAGPAEPPEASGG---DLELLEPELDCGQDWDLDEVMNMLVLDSPEEDVAISLLAPAPD
scaIRF3         ------------------------------------------------EVPGGPGQPPGAGAAGPEEPPEXAGG---DLELLEPELDCGQDWDLDEVMNMLVLSSPEEDXAISLLAPAPD
rpeIRF3         ---------------------------------------------------EAPGQPPGAGGAGPEMPPEVPGGDLEDLELLEPELDCGQDWDLDEVMNMLVLSSPDENAAISLLGPAPP
ccarIRF3        -------------------------------------------------NGARAGADEAPSAAVGGTEADASSQ---LLDRSGGGLSCSQDDDLDDILNALALSSPEEDAPCSALA---P
asiIRF3         -------------------------------------------------VGVQAGADEAPLVAGRADTLDGPLE---QLLGEG----CSPDSDLEQYVHLMTLCSPEEDISSPDLS---L
hsaIRF3         ----------------------------------------------------SGVGDFSQPDTSPDTNGGGSTS-------------DTQEDILDELLGNMVLA-PLPDPGPPSLA----
cfaIRF3         ----------------------------------------------------SGAGNLPELDTFPDTNGRYSTS-------------DTQEDTLEELLGDMVLT-PFPDEGPSSLV----
mmuIRF3         ----------------------------------------------------PGARDFVHLGASPDTNGKSSLP-------------HSQEN-LPKLFDGLILG-PLKDEGSSDLA----
                                                                              :         .                  :  * . .  : *   *   :    . *

dnoIRF3         DAPDPPVLPPDPPYCPPEDAMGLGPPPQP-GCPLP-DLLIGADELKTDLEVRVYYRGRAVLQTTVAAPGQ-LLLVAPGTPGTPV-APGTVALPEPA--VADGRQARYTRRVLRGLGAGVG
scaIRF3         AAPDPSLLLPDPPYCPPEDAVGLGPPLQP-SSPLP-DLLIGADELRTDLEVRVYYRGRPVLQAAVAAPGE-LRLVPP--PGPPG-AAGTVALPEPA--VADGRQARYTRRVLQGLGAGVG
rpeIRF3         AAPGPPALLPDPPYCPPEEAVGPPPALQP-SSPLP-ELLIGADELRTDLEVRVYYRGRPVLQATVAAAPEGLRLAAPGPAASAG-VGAAVPLPEPE--VPDGRQARYTRRVLQGLGAGVA
ccarIRF3        GAP--YVGDAATGGALSPGFTDFPPLVQP--APSALEQLLGSNVLMTDFEVRVYYRGHLVLQTLVPNPQG-FRLVSTSPGPSPCPELVDVELPGPGM-LPDRVQASYTARLLQGLGAGVL
asiIRF3         FGPGEAVGDAAVP-VLS-GAEVLPPLVQPLSTPNPGGLLLGSNEL--DFDVNVYYRGHRVLHTLVSSSQG-LRLVFPQSSPSPAPELTDVVLPDPES-LPDRVQANYTGRLLRGLGPGVV
hsaIRF3         -----VAPEPCPQPLRSPSLDNPTPFPNLGPSENPLKRLLVPGEE-WEFEVTAFYRGRQVFQQTISCPEG-LRLVGSEVGDRTLPGWP-VTLPDPGMSLTDRGVMSYVRHVLSCLGGGLA
cfaIRF3         -----VVPEQTPPLLLSPTIDLPAPCPNSEPPENPLKRLLVPDEE-WEFEVTAFYRGRQVFQQTVSCPRG-LRLVAAAGGDTMLPGQP-IILPDPGVLVTDKTVMGYVRRVLSCLGGGLA
mmuIRF3         -----IVSDPS-QQLPSPNVNN---FLNPAPQENPLKQLLA-EEQ-WEFEVTAFYRGRQVFQQTLFCPGG-LRLVGSTA-DMTLPWQP-VTLPDPEGFLTDKLVKEYVGQVLKGLGNGLA
                                .           :       .     *:        :::* .:***: *::   :    .    : *.  .                : ** *     :.*        *.  ::*    ** *:

dnoIRF3         LRARGPGLEGARLGCCRVFWGRGGT--PGPGP-PRGALPKGTYGPLYGLRDFVRELIAFMEGGGESPQYELWLCLGEPWPEPGHSWTRKLIMVQVVPVALRLLHELSRAQGASSLRSSEL
scaIRF3         LRARGPALEGTRLGRCRVFWGRGGG--PGPGA-PGGALPKGTYGPLYGLRDFVRELIAFLEGRGGSPHYELWLCLGEPWPEPSHSWTRKLVMVQVVPVALRLLHELSQEQGASSLRSSEL
rpeIRF3         LRARGPELEGARLGRCRVFWGRGGGGNPGPGA-PGGALPKGTYGPLYGLRDFVRELIAFMEGRGGHPXYELWLCLGESWPEPGHHWTRKLVMVQVVPVALRLLHELSRAQGASSLRSSEL
ccarIRF3        VRVEGPALCATRLGRLHAYWGHTET--PEPGA-EGGELSKEGYTRLYDMSHFVRELICFIEGQGSSPTYELWLCLGEAWPDTGRSWKKKLIMVQVVPTVLRTLHELGQAGGASSLRGEEL
asiIRF3         LRAKGRILGGCRLGRCHAFWGLTAT--PAPGA-LGGEIPKESFTPLYSLPEFIQDLIGFMEGQRRSPDYELWLCLGEKWPDTERPWNRKLIMVQVVLVALQKLHELSQAGGASSLRSSEL
hsaIRF3         LWRAGQWLWAQRLGHCHTYWAVSEELLPNSGHGPDGEVPKDKEGGVFDLGPFIVDLITFTEGSGRSPRYALWFCVGESWPQ-DQPWTKRLVMVKVVPTCLRALVEMARVGGASSLEN-TV
cfaIRF3         LWRAGQQLWARRLGHCHTYWALGEELLPDSSPRPGGEVPKDEDGDLFDLRPFVSDLIAFIKGSRHSPRYTLWFCVGEPWPQ-DQPWTKKLVMLKVVPTCLRALLEMARLEGASSLEN-TV
mmuIRF3         LWQAGQCLWAQRLGHSHAFWALGEELLPDSGRGPDGEVHKDKDGAVFDLRPFVADLIAFMEGSGHSPRYTLWFCMGEMWPQ-DQPWVKRLVMVKVVPTCLKELLEMAREGGASSLK--TV
                :    *  *  . ***  :.:*.        *  ..     *  :  *       ::.:   *:  :**  *  :*     *  *  **:*:**  **:   :  *  ::*:*::**  .  *:  *  *:.:   *****.    :

dnoIRF3         QLQLSASLG-----EGGLLGALRSWDERMDSLPSA--
scaIRF3         QLQLSASLG-----PGGLLGALRGWGERMDTLPXA--
rpeIRF3         QLQLSASLDGADAGGGGLLGALRSWDERMDSRP----
ccarIRF3        DLRISDSLG-----EAGLLGALRDWEERMESQPYS--
asiIRF3         NLHISDSLG-----DGGLLGALRSWEELMDTQP----
hsaIRF3         DLHISNSHP-LSLTSDQYKAYLQDLVEGMDFQGPGES
cfaIRF3         DLHISNSYP-LTLTSDQYKAYLQDLVEDMDFWVTGEV
mmuIRF3         DLHISNSQP-ISLTSDQYKAYLQDLVEDMDFQATGNI
                :*::* *               . *:.  * *:
```

**Suppl. Fig. 1** CLUSTAL 2.1 multiple sequence alignment of avian IRF3 proteins (dno – *Dromaius novaehollandie*, sca – *Struthio camelus*, rpe – *Rhea pennata*) with proteins from non-avian archelosaurs (ccar – *Caretta caretta*, asi – *Alligator sinensis*) and mammals (hsa – *Homo sapiens*, mmu – *Mus* *musculus*, cfa – *Canis familiaris*). Exonic structures of CDS are shown by alternating black and blue ink with amino acids which codons are split by introns shown in red ink. Five conserved tryptophans in DBD are shown on cyan highlight while two conserved serines in the C-terminus instrumental for activation of IRF3 by phosphorylation are on yellow highlight. Four-fold repetitive structure of the extension of the second exon of emu (dno) IRF3 is visualized using alternating single and double underlines.

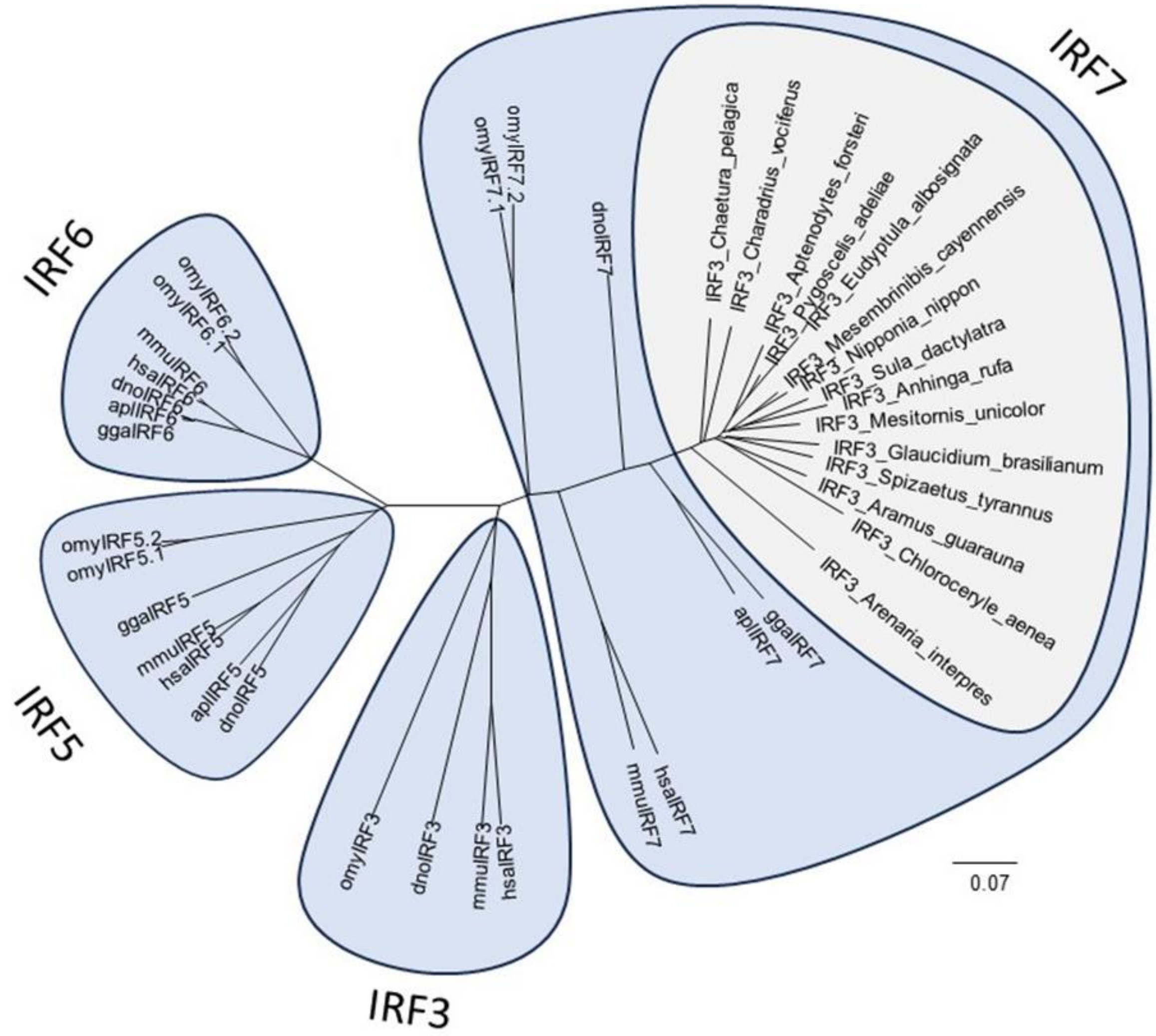


**Suppl. Fig. 2** IRF7 sequences of neognath birds that are erroneously annotated as IRF3 in genebank (shown on gray background) cluster with vertebrate IRF7 sequences rather than with IRF3. NJ tree of Clustal X-aligned protein sequences. False IRF3 sequences are listed in Supplementary table 1. Control IRF sequences come from following species: apl - Anas platyrhynchos (duck), dno - Dromaius novaehollandie (emu), gga - Gallus gallus (chicken), hsa - Homo sapiens (human), mmu - Mus musculus (mouse), omy - Oncorhynchus mykiss (rainbow trout) and are available in Suppl. Tab. 5.

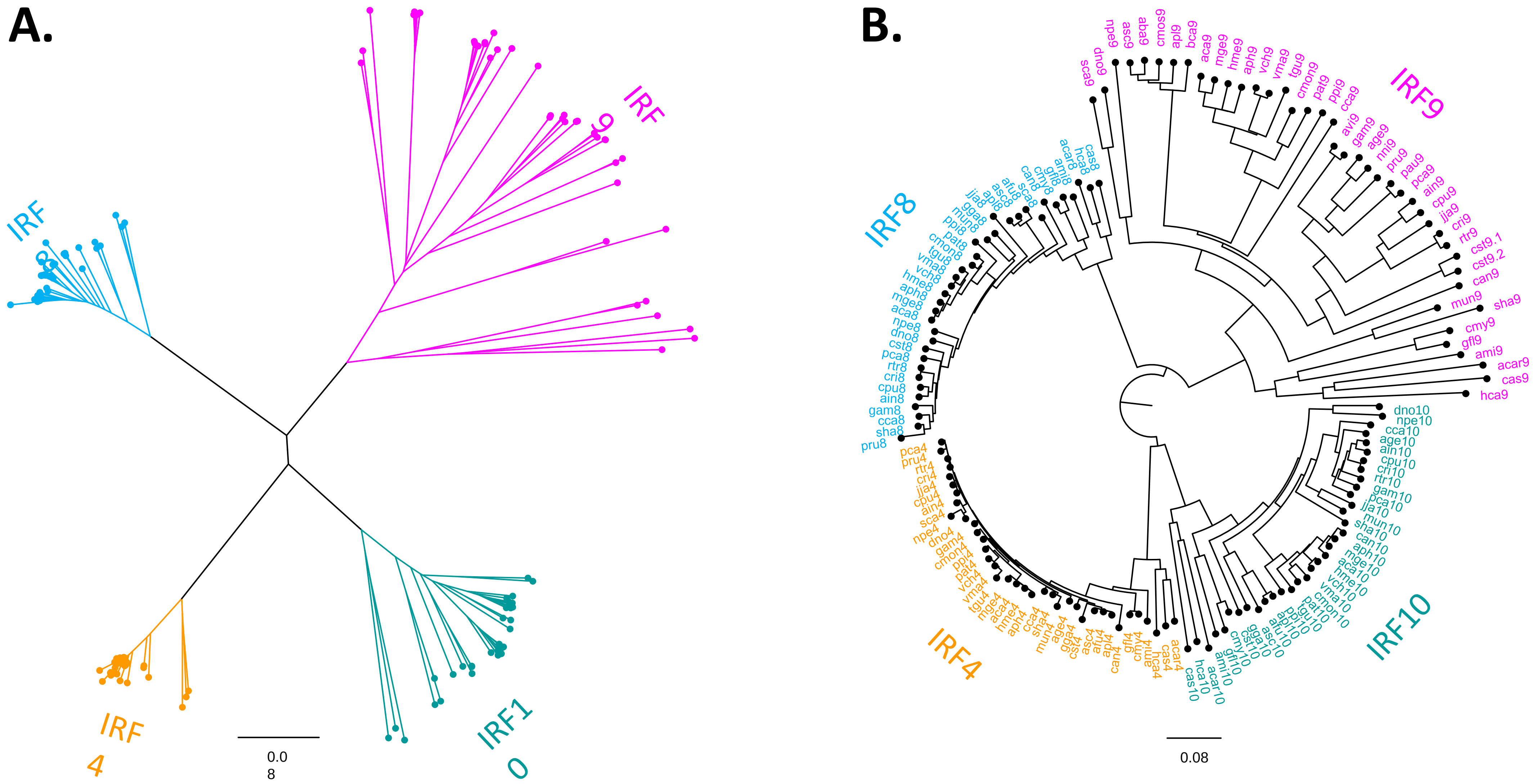


**Suppl. Fig. 3** Supplementary information for Fig. 4. Avian IRF9 protein sequences form a single clade in the phylogenetic tree. **A.** The tree shown in the same form of radial phylogram as is shown in Fig.4 in the main text. Each IRF family member is, however, distinctly color-coded. **B.** The same tree shown in form of circular cladogram with species lables included. For species labels look the Suppl. Tab. 5.

```
duck IRF9    1 ATGACGTCAGGGCGGCGCCGCCTCCTGCCCTGGCTGCTGGCGCAGGCGCAGAGCGGCCGCTTCCCGGGGCTGGAGTTCGACGACGCCGCCCGCAGCGCCCTCAGGG  106
synthesis    1 ATGACCAGCGGAAGAAGGCGCCTGCTGCCTTGGCTGCTGGCTCAGGCACAGAGCGGAAGATTCCCAGGGCTGGAGTTTGATGACGCAGCTAGGTCCGCACTGCGCG  106

duck IRF9  107 TGCCCTGGGAGCGCGCCGGAAGGGGCGGGGCCGGGGGCGGGGCCGCCGCCGCCCTCTGCCAGGCGTGGGCGGAGTACAAAGGGGCCACGCCCCCGCCGGGCCCGGC  212
synthesis  107 TGCCTTGGGAAAGGGCAGGACGCGGCGGAGCCGGGGGCGGAGCAGCAGCTGCACTGTGCCAGGCATGGGCTGAGTACAAGGGAGCAACTCCACCTCCAGGACCAGC  212

duck IRF9  213 CGTCTGCAAGACCCGGCTGCGCTGCGCCCTGCACAAGAGCCCCGAGCTGCAGGAGGTGCCCGAGCGCGCCCGCCTGGACGGGCCCCGCCCCTACAAGGTGTACCGC  318
synthesis  213 TGTGTGCAAGACCAGGCTGAGATGCGCCCTGCACAAAAGCCCAGAACTGCAGGAAGTGCCCGAAAGGGCTAGGCTGGATGGACCAAGGCCTTACAAAGTGTACAGA  318

duck IRF9  319 CTGCTGGGGCCACGCCCACCACGCAACGGCCCCGCCCCCGGCGCGGCTCCAAGGGGCGCAAGGAGGAGGCGGAGCCACGCGGCGATGACGCGACCCCGCCCCCCA  424
synthesis  319 CTGCTGGGACCAAGGCCACCTAGAAACGGACCAGCACCACGGAGAGGAAGCAAGGGCAGGAAAGAGGAAGCAGAGCCTCGCGGAGATGACGCAACACCACCACCTA  424

duck IRF9  425 GCTCTGAGGAGGACGAGGAGGAGGCGGAGCCGAGCCCAGCCCCGCCCCCTCCCGCTGCTGTGACCCCGCCCCCTTCCCGCCAGCTCGCCATCCTATTGGACAGCCC  530
synthesis  425 GCTCCGAGGAAGACGAGGAAGAGGCTGAACCTTCCCCAGCACCACCACCTCCAGCAGCTGTGACTCCACCTCCAAGCAGACAGCTGGCTATCCTGCTGGATTCCCC  530

duck IRF9  531 GGAGCCCCTCCCCCCAGCCGAAGGCGACGCGGCATTGGGCGTCTCCCTCTGGGTGGGCGGGGCCCTGGCCTGGAGGGCGTGGCTCCCGGCGGGCGAGTACCTCCTA  636
synthesis  531 AGAGCCTCTGCCACCTGCAGAAGGAGACGCAGCACTGGGAGTGAGCCTGTGGGTGGGGGGCGCACTGGCATGGAGGGCATGGCTGCCAGCAGGAGAGTACCTGCTG  636

duck IRF9  637 TTGGCCGCCGGCAGCCCCGCCCCCGGCCCCGCCCCCTTGCCCCGCCTCCTGGTGCCGGCCCCGCCCCCCCCGGCCCTGGAGGCGCTGGGGCGGGGCTGGCCGGG  742
synthesis  637 CTGGCTGCAGGATCCCCAGCTCCAGGACCTGCACCACTGCCAAGACTGCTGGTGCCCGCTCCTCCACCTCCAGCACTGGAAGCACTGGGAGCTGGACTGGGAAGGG  742

duck IRF9  743 GGCTGCTGGTGGCCAGCGGGGGGCGGGGCCTCTTCCTCAGGACACGCCCCCGGGGGCGGGGCCTCGCCCTCAGCGCCCCCCACGCCGGGGAGCCGGGGGGCGGGGC  848
synthesis  743 GACTGCTGGTCGCCAGCGGAGGGAGAGGACTGTTCCTGAGGACAAGACCCAGGGGAAGGGGACTGGCACTGTCCGCCCCTCACGCTGGAGAGCCAGGCGGAGGGGC  848

duck IRF9  849 CTGGCCACAGGGGGAGGAGCCACTGGTGGCCGCCTTCGACAGCCGGCGCTTCCGGGAGGAGCTGCAGCGGCACCGCCAGGGGCTGGGCCCCCCCCCGAGCACCGC  954
synthesis  849 ATGGCCCCAGGGCGAAGAGCCTCTGGTGGCCGCTTTCGATAGCAGGCGCTTTAGGGAAGAGCTGCAGAGGCACAGACAGGGACTGGGACCACCTCCAGAGCACAGG  954

duck IRF9  955 GTGCTGCTGGCGGCCACCCCCGGGGAGCTCCGCACGGGGGCGGGGCCAGCGCACAGGGGGCGGTGCTACAGCTGGAGCAGGCCCTGGCCCAGCGCCTCCTGGGCC 1060
synthesis  955 GTGCTGCTGGCAGCAACCCCAGGAGAACTGAGGACAGGCGGAGGGGCTTCCGCACAGGGAGCTGTGCTGCAGCTGGAGCAGGCACTGGCTCAGAGGCTGCTGGGAC 1060

duck IRF9 1061 CCGCGCCGGATCCCGCCAATTAA  1083
synthesis 1061 CTGCACCAGACCCAGCTAATTGA  1083
```

**Suppl. Fig. 4** Pairwise alignment of IRF9 coding sequences of endogenous duck IRF9 with in vitro synthesized construct. The predicted protein coding sequence is identical for both. In the gene synthesis construct, the codons were optimized to break GC-rich stretches. Mismatches in the nucleotide sequence are highlighted in gray color.
